# Plant Cleavage Factor I complex is essential for precise cleavage and polyadenylation site determination

**DOI:** 10.1101/2024.03.28.587165

**Authors:** Lukasz Szewc, Xiaojuan Zhang, Mateusz Bajczyk, Dawid Bielewicz, Marta Zimna, Kei Yura, Mariko Kato, Mika Nomoto, Marta Garcia-León, Vicente Rubio, Yasuomi Tada, Tsuyoshi Furumoto, Takashi Aoyama, Zofia Szweykowska-Kulinska, Dorothee Staiger, Artur Jarmolowski, Tomohiko Tsuge

## Abstract

Cleavage factor I (CFI) is a four-subunit protein complex of the pre-mRNA 3’ end processing machinery in eukaryotes. In Arabidopsis, AtCFI25a, AtCFI25b, AtCFI59, and AtCFI68 have been identified as potential components of AtCFI, *in silico*. Here, we show that the AtCFI25a, AtCFI59, and AtCFI68 proteins each pulled down all components of the CFI, confirming that these subunits form the plant CFI complex. Furthermore, either AtCFI59 or AtCFI68 was essential for nuclear localization of the smallest subunit, AtCFI25a. Mutants with single loss-of-function for *AtCFI59* or *AtCFI68* showed no obvious morphological defects compared to wild-type plants, while the double mutant displayed pleiotropic morphological defects, identical to those previously reported for *AtCFI25a* loss-of-function plants. Moreover, these morphological defects correlated with alterations in the usage of 3’ UTR cleavage and polyadenylation sites. *atcfi25a*, *atcfi25a atcfi25b* and *atcfi59 atcfi68* double mutants showed widespread changes in the choice of cleavage and polyadenylation sites. In most cases, more proximal cleavage and polyadenylation sites were used, leading to shorter 3’ UTRs. In particular, genes involved in light intensity, light harvesting, photosynthesis and cold responses showed significant dependence on AtCFI function. Furthermore, transcripts coding for AtCFI subunits showed altered 3’ end processing in these mutants, suggesting self-regulation function of AtCFI in plants.

## INTRODUCTION

One of the key steps in the biogenesis of functional eukaryotic messenger RNA is the formation of its 3’ end. This process is called <u>c</u>leavage and <u>p</u>oly<u>a</u>denylation (CPA). CPA starts with the cleavage of a precursor mRNA (pre-mRNA) molecule and ends with the template-independent addition of series of adenine nucleotides (A) to the 3’ end of mRNA. The poly(A) tail serves important functions: It protects mature mRNAs from degradation [Saini *et al*. 1990, Manguys *et al*. 2003, Wigington *et al*. 2014, Park *et al*. 2023], promotes their export to the cytoplasm [Manguys *et al*. 2003, Apponi *et al*. 2010, Iglesias *et al*. 2010], helps with the prevention of nuclear export of unspliced pre-mRNAs [Kwiatek *et al*. 2023], and enhances translation of mRNAs in the cytoplasm [Manguys *et al*. 2003, Kühn and Wahle 2004, Wigington *et al*. 2014, Lima *et al*. 2017, Passmore and Tang 2021].

Most of the pre-mRNA possess diverse lengths of untranslated regions at its 3’ end (3’ UTR). Genome-wide analyses of polyadenylation sites in *Arabidopsis thaliana* reported that 70% or more of all genes possess two or more poly(A) site clusters [Wu *et al*. 2011]. The selection of the polyadenylation site involves a complex interplay between polyadenylation signal sequences surrounding the CPA site, as well as pre-mRNA 3’ end processing protein complexes which recognize these motifs [Zhang *et al*. 2021, Rodríguez-Molina and Turtola 2023]. The plant polyadenylation signal consists of three weakly conserved sequence regions: Far Upstream Element (FUE), Near Upstream Element (NUE), and Cleavage Element (CE) [Hunt 2008, Xing and Li 2011]. The FUE is located about 30-50 nt upstream the CPA site and is rich in U>A>G [Hunt 2008, Xing and Li 2011]. The NUE is located approximately 10 to 40 nt upstream from the CPA site and is A-rich. The NUE is an equivalent of the highly conserved canonical mammalian polyadenylation signal, the A(A/U)UAAA motif [Proudfoot and Brownlee 1976], however, such sequence is found in less than 10% of the studied *A. thaliana* genes [Loke *et al*. 2005]. Interestingly, recent studies show that a similar but shorter sequence, the UAAA motif, is commonly present upstream of plant CPA sites [Ye *et al*. 2021]. The CE, consists of the cleavage site (UA or CA dinucleotide) surrounded by an U-rich region (U>A>C rich) [Xing and Li 2011]. Although there is no sequence conservation within *cis*-elements located in the 3’ ends of pre-mRNA between vertebrates, plants, and yeasts, the general architecture of polyadenylation signals across kingdoms seems to be very similar [Millevoi and Vagner 2010, Bernardes and Menossi 2020].

This similarity also applies to the composition of the multiprotein complexes that recognize the polyadenylation signals. Higher plants share with yeast and/or mammals the core components involved in 3’ end processing that can be divided by function into four main subcomplexes, also referred to as factors. The first subcomplex, Cleavage and Polyadenylation Specificity Factor (CPSF), consists of six main proteins: CPSF160, CPSF100, CPSF73, CPSF30, FY (ortholog of the mammalian WDR33 protein), and AtFIP1. The second subcomplex, Cleavage Stimulatory Factor (CstF), is composed of CstF64, CstF77, and CstF50, although CstF50 can be absent in some plants [Hunt *et al*. 2012]. The third subcomplex, Cleavage Factor II (CFII), is comprised of PCF11 and CLP1 in yeast and mammals. Interestingly, in plants these proteins are usually encoded by more than one gene (in *A. thaliana*, PCF11 is encoded by at least two genes and CLP1 is encoded by two genes) [Hunt *et al*. 2008, Hunt *et al*. 2012]. The fourth subcomplex is Cleavage Factor I (CFI). Its presence in plants has been suggested based on computational studies [Hunt *et al*. 2012], until its physiological role was recently confirmed through experiments [Zhang *et al*. 2022]. Mammalian CFI (CFIm) has been identified as one of the key regulators of alternative polyadenylation, the process which generates distinct 3’ ends of mRNA originated from the same gene [Hardy and Norbury 2016]. Many eukaryotic genes possess more than one cleavage and polyadenylation site [Tian *et al*. 2005, Zhang *et al*. 2021], and the choice of CPA site introduces another layer of regulation of gene expression. Selection of CPA site can influence the stability of mature mRNA [Hogg and Goff 2010, Hoffman *et al*. 2016], mRNA nuclear export [Chen and Carmichael 2009], its cellular localization [An *et al*. 2008], cellular localization of translated protein [Reid and Nicchitta 2015], and production of distinct protein isoforms [Alt *et al*. 1980].

The human CFIm complex forms a tetramer, composed of two small CFIm25 subunits, forming a homodimer, and two larger subunits of CFIm59 and/or CFIm68. Each of the CFIm25 subunits can interact with one CFIm59 or CFIm68 protein [Kim *et al*. 2010, Gruber *et al*. 2012]. In humans CFIm regulates poly(A) site selection through the recognition of UGUA sequence by CFIm25 [Yang *et al*. 2010], but the interaction of CFIm25 with CFIm59 and/or CFIm68 enhances its RNA binding affinity [Yang *et al*. 2011]. Despite the fact that CFIm can be formed by two or three proteins, many studies have suggested that the alternative polyadenylation activity of CFIm is mediated by CFIm25 and CFIm68 only. Knockdown of CFIm25 and CFIm68, but not CFIm59, lead to reduced usage of distal poly(A) sites and general 3’ UTR shortening [Gruber *et al*. 2012, Martin *et al*. 2012, Zhu *et al*. 2018, Li *et al*. 2023]. Recent data show that this alteration at the 3’ UTR is caused by a distinct preference of CFIm68 and CFIm59 for specific subset of UGUA elements. Although, CFIm68 and CFIm59 can partially compensate for each other in the CFIm complexes across all alternative polyadenylated genes, CFIm68 is found to be specifically enriched upstream of distal cleavage sites, thus mRNAs are shortened upon CFIm68 depletion [Tseng *et al*. 2022].

In this report we present the composition and function of the plant CFI complex in *A. thaliana*, which consists of a small subunit AtCFI25a (At4g25550) and two larger subunits: AtCFI59 (At1g13190) and/or AtCFI68 (At5g55670). We demonstrated that these proteins form a complex in the plant cell nucleus, most likely through direct interaction of AtCFI25a with AtCFI59 and/or AtCFI68. Furthermore, direct interactions between AtCFI59 and AtCFI68 subunits were described, suggesting that the two larger subunits can form homo- and heterodimers. *atcfi25a*, *atcfi25a atcfi25b* and *atcfi59 atcfi68* double mutants showed widespread changes in the choice of CPA sites. In most cases, more proximal CPA sites were used, leading to longer 3’ UTRs. Taken together, these results shed light on the molecular mechanism of AtCFI-regulated CPA site choices at the global gene level.

## MATERIAL AND METHODS

### Plant material and growth conditions

*Arabidopsis thaliana* (L.) Heynh. Columbia (Col) ecotype was used as wild-type plants. T-DNA insertion lines of interest, SALK_090357 (*atcfi59-1*), SALK_128104 (*atcfi59-2*), GABI_217C05 (*atcfi68-1*), GABI_213F12 (*atcfi68-2*), SALK_036546 (*atcfi68-3*), were identified and obtained accordingly in the collection of the Salk Institute Genomic Analysis Laboratory (http://signal.salk.edu) [Alonso *et al*. 2003], and the GABI-Kat collection (https://www.gabi-kat.de) [Kleinboelting *et al*. 2012]. Homozygous lines were established through genotyping using primer sets shown in Supplementary Table S1 in respect to T-DNA insertion. Formally characterized null mutant alleles for *AtCFI25a* and *AtCFI25b* [Zhang *et al*. 2022] were utilized accordingly for corresponding experiments. Double mutants, *atcfi25a atcfi25b* and *atcfi59 atcfi68,* were generated by respective crosses. Seeds were surface sterilized and imbibed at 4°C for 3 days, before plating on full or half strength Murashige and Skoog (MS) medium (0.8% agar, B5 vitamins, 2.5 mM MES, and 1% sucrose, pH 5.7) or sowing on soil. Plants were grown under continuous light or 16h light / 8h darkness cycle, at 22°C.

### DNA constructs and plant transformation

For *AtCFI59_prom_::GUS* and *AtCFI68_prom_::GUS* constructs, DNA fragments encompassing 1428 and 933 bps intergenic sequences, upstream from the start codons, were amplified by PCR using primers listed in Supplementary Table S1 and genomic DNA isolated by ISOPLANT (Nippon Gene, Tokyo, Japan). Fragments were cloned into cloning vector pPCR-Script and then to binary vector PBI101 using *Bam*HI, *Not*I, *Sal*I, and corresponding blunt-ended sites. Binary vectors were introduced by electroporation into *Agrobacterium tumefaciens* LBA4404 or AGL-1, that was then used to transform *A. thaliana* by the floral dip method [Clough and Bent 1998]. The resulting transgenic plants were self-pollinated, and T3 plants homozygous for the transgene were used in subsequent experiments.

Full-length coding sequence of *AtCFI59* and *AtCFI68* was amplified by PCR using primers listed in Supplementary Table S1, and cDNA mixture prepared from total seedling RNA using SuperScript^TM^ III First-Strand Synthesis System (Invitrogen/Thermo Fisher Scientific, Waltham, USA). Each fragment was cloned into pPCR-Script vector and further cloned into suitable expression vectors for corresponding experiments.

For constructs designed to express proteins fused to the tags used in the pull-down assay, coding sequences of *AtCFI25a* and *AtCFI59* were cloned into pGEX5X-2 (GST) (Merck/GE Healthcare, Darmstadt, Germany) or pET30c (HIS) (Merck/Novagen, Darmstadt, Germany) using *EcoR*I and *Not*I sites. The constructs were then transformed into *Escherichia coli* BL21 (DE3) to express recombinant protein. For co-immunoprecipitation (co-IP) experiments, coding sequences of *AtCFI25a*, *AtCFI59*, and *AtCFI68* were amplified from cDNA templates, and cloned into pUBN-GFP-Dest and pUBC-GFP-Dest vectors [Grefen *et al*. 2010] to be expressed in *atcfi25a*, *atcfi59*, and *atcfi68* mutants. For FRET-FLIM experiments, corresponding coding sequences were cloned into modified pSAT6-DEST-EGFP vector [Tzfira *et al*. 2005]. In this vector the 35S Cauliflower Mosaic Virus (CaMV) promoter was exchanged by the *A. thaliana* ubiquitin-10 promoter. To obtain tagRFP-fusion proteins, the eGFP coding sequence was replaced by the tagRFP [Merzlyak *et al*. 2007] coding sequence.

For mutated NLS versions of AtCFI59 and AtCFI68 proteins, shorter coding sequences, without the part containing the NLS, were cloned into pENTR vector and then cloned *via* Gateway recombination cloning to pSAT6-DEST vectors.

All constructs using PCR in its cloning processes were sequenced to confirm correct sequences.

### β-Glucuronidase (GUS) staining

Plant material at different stages was collected, fixed, and stained for GUS activity [Imajuku *et al*. 2001]. X-Gluc was used for histochemical staining to monitor GUS activity. Multiple independent lines, homozygous for the transgene, were examined.

### RT-PCR analysis

Total RNA was isolated from different organs of *A. thaliana* using ISOGEN-kit (Nippon Gene, Tokyo, Japan). Growth stages of samples are noted in each experiment and figure legend. For semi-quantitative analyses, SuperScript™ III One-Step RT-PCR System with Platinum™ Taq DNA Polymerase (nvitrogen/Thermo Fisher Scientific, Waltham, USA) were used to detect transcripts of *AtCFI59* and *AtCFI68* with primer sets shown in Supplementary Table S1. PCR products were analyzed by agarose gel electrophoresis.

### Pollen observations using Alexander’s staining method

Pollen grains were mounted on slides in Alexander’s staining solution [Alexander 1969] and examined under the microscope (MZFL III Leica Microsystems, Wetzlar, Germany). Viable and non-viable pollen grains were observed in three independent experiments.

### *In vitro* pull-down assay

Recombinant proteins were expressed in *E. coli* and prepared in hypotonic buffer (pH 7.2, 20 mM HEPES pH 7.0, 1.5 mM MgCl_2_, 5 mM KCl, 1× protease inhibitor cocktail (Nacalai Tesque, Kyoto, Japan) by sonication. GST-tagged AtCFI59 was purified using agarose beads (MagneGST^TM^ Glutathione Particles; Promega, Madison, USA) and retained on the beads to pull-down purified HIS-tagged AtCFI25a protein in TBS buffer (50 mM Tris-HCl pH 7.4, 150 mM NaCl, 1 mM DTT, 1 mM PMSF, 50 µM MG132, 1× protease inhibitor cocktail (Nacalai Tesque, Kyoto, Japan). After washing the beads three times with TBS buffer with 0.5% Triton X-100, the HIS-tagged AtCFI25a protein bound to GST-AtCFI59 was detected by western blot analysis using an anti-HIS antibody (G18) (sc-804; Santa Cruz Biotechnology, Dallas, USA) at 1:5000 dilution.

### Co-immunoprecipitation experiments and mass-spectrometry analyses

For co-immunoprecipitation (co-IP), seeds were stratified on solid half strength MS medium for 3 days at 4°C and transferred to 22°C under the 16h light / 8h darkness cycle. Seedlings were grown for 16 days (∼14 days after germination), harvested and frozen in liquid nitrogen. For each co-IP, 0.6 g of plant material ground in liquid nitrogen was extracted with 1.5 ml of lysis buffer (50 mM Tris-HCl pH 8.0, 75 mM NaCl, 1% Triton X-100, 2x cOmplete™ EDTA-free protease inhibitor (Roche, Mannheim, Germany), 10 µM MG132 (Sigma-Aldrich, Damstadt, Germany)). Cell debris was removed by two rounds of centrifugation (10 and 5 min, 16,000xg at 4°C). Supernatants were incubated for 30 min with anti-GFP antibodies coupled to magnetic beads (μMACS GFP isolation Kit; Miltenyi, Bergisch Gladbach, Germany). Supernatants with beads were loaded on μMACS separation columns (conditioned with lysis buffer) and washed four times with 300 µl of washing buffer (50 mM Tris-HCl pH 7.5, 0.1% Triton X-100). Samples were eluted with 100 µl of pre-warmed (95°C) elution buffer (µMACS GFP isolation Kit, Miltenyi, Bergisch Gladbach, Germany). Control IPs were performed on wild-type plant extracts with anti-GFP antibodies. Eluted immunoprecipitated protein was precipitated by adding TCA and Tween 20 (Sigma Aldrich, St. Louis, USA) to a final concentration of 10% and 0.5% respectively and incubated on ice for 30 min. Samples where then centrifuged for 10 min at 14,000 rpm at 4°C. Pellet was washed once with cold 10% TCA and twice with cold 90% acetone. Each wash was followed by the pellet centrifugation for 10 min at 14,000 rpm 20,000xg at 4°C. After the last centrifugation, the supernatant was removed, and the pellet was air-dried at room temperature.

Mass spectrometry analysis was performed in the Mass Spectrometry Laboratory (Institute of Biochemistry and Biophysics, Polish Academy of Science, Warsaw, Poland). Purified proteins were digested with sequencing-grade trypsin (Promega, Madison, Wisconsin, USA) and separated by nanoAcquity UPLC (Waters, Milford, Massachusetts, USA) liquid chromatographs connected to Orbitrap Elite (Thermo, Waltham, Massachusetts, USA) for AtCFI25a co-IP and AtCFI68 co-IP elutes, or Evosep One (Evosep Biosystems,Odense, Denmark) connected to Exploris 480 (Thermo, Waltham, Massachusetts, USA) for AtCFI59 co-IP elute. Data were pre-processed with Mascot Distiller (Matrix Science, London, UK) and searched against the TAIR10 database with Mascot Server (Matrix Science, London, UK). The total number of MS/MS fragmentation spectra was used to quantify each protein in all replicates. The statistical analysis based on spectral counts was performed using the IPinquiry R package [Kuhn *et al*. 2023] that calculated fold change and p values using the quasi-likelihood negative binomial generalized log-linear model implemented in the edgeR package [Robinson *et al*. 2010]. The size factor used to scale samples was calculated according to the DESeq2 normalization method [Love *et al*. 2014]. P values were adjusted using the Benjamini– Hochberg method from the stats R package.

### Förster resonance energy transfer (FRET) - fluorescence lifetime imaging microscopy (FLIM) analysis

Protoplasts were prepared as previously described [Bajczyk *et al*. 2020]. Protoplasts were incubated in the dark for 10-12 h at 22°C before FRET-FLIM analyses. FRET-FLIM analyses was performed as previously described [Bajczyk *et al*. 2020]. For each sample, more than twenty cells from at least three biological repeats, with independent protoplast isolation and transformation, were used. The results were evaluated with the two-sided Mann-Whitney-Wilcoxon test (p-value < 0.001).

### Rapid amplification of cDNA ends (RACE)

Total RNA was isolated from 7 days old seedlings of *A. thaliana* using ISOGEN (Nippon Gene Co, Toyama, Japan). 3’ RACE was performed using 3’-Full RACE Core Set (TaKaRa, Kusatsu, Japan). PCR amplification on the 3’ UTR regions of *AtCFI25a*, *AtCFI25b*, *AtCFI59*, and *AtCFI68* transcripts were carried out using nested primer sets shown in Supplementary Table S1. PCR products were analyzed by agarose gel electrophoresis.

### PAT-seq experiments and analysis

For poly(A) tag sequencing (PAT-seq) [Pati *et al*. 2015] experiments, seeds were stratified on solid half strength MS medium for 3 days at 4°C and transferred to 22°C under 16h light / 8h darkness cycle. Seedlings were harvested at 14 days after germination and frozen in liquid nitrogen, to be stored at −80°C until use.

For each RNA isolation, 600 mg of plant material ground in liquid nitrogen was utilized. Total RNA was extracted with RNeasy Plant Mini Kit (Qiagen, Hilden, Germany) and checked for quality with Agilent RNA 6000 Nano Kit (Agilent Technologies, Waldbronn, Germany). Only RNA samples with RNA integrity number (RIN) > 9 were used for further steps. 25 µg of total RNA was then used for zinc-mediated RNA fragmentation [Pati *et al*. 2015] and purified with RNeasy Mini Kit (Qiagen, Hilden, Germany). Polyadenylated RNA fragments (150 – 500 nt) were enriched with Dynabeads™ mRNA Purification Kit (Thermo Fisher, Vilnus, Lithuania) and used with the mixture of specially designed primers (SWITCH2 and RT_universal_primer, Supplementary Table S2) for cDNA synthesis (SMARTScribe Reverse Transcriptase, TaKaRa, Kusatsu, Japan).

In the following step, RNA was removed from the mixture with RNAase H and RNase A/T1. cDNA fragments were purified with AMPure XP (1.8 of sample volume, Beckman Coulter, Brea, USA). Libraries were amplified with Phire Hot Start II DNA Polymerase (Thermo Fisher, Vilnus, Lithuania) and barcode PCR primers (NEXTflex™ Small RNA-Seq Kit v3; Austin, USA). After amplification libraries were purified with 0.8 volume of AMPure XP beads and tested on Agilent DNA 1000 or Agilent HS dsDNA Chips. Sequencing was performed on the NextSeq Illumina platform (1×75 with 20% of Phix spike) at Fasteris in Switzerland.

From the raw reads, the first six nucleotides (NNNNNN - unique molecular identifiers (UMI)) were removed by the UMI tool function [Smith *et al*. 2017], and then the next 19 nucleotides were trimmed with the cutadapt tool (https://cutadapt.readthedocs.io/en/stable/installation.html#installation-on-debian-ubuntu) [Martin 2011]. In case the first present nucleotide was a “T”, this was also removed. Clean reads were reverse complemented with the fastx_reverse_complement tool, and then aligned to the Arabidopsis TAIR10 reference genome with Hisat2 software [Kim *et al*. 2019] using the “--no-spliced-alignment” option. QuantifyPoly(A) tool [Ye *et al*. 2021] was used to analyze polyadenylation sites. In order to do that, Binary Alignment/Map (BAM) files from the hisat2 alignment were converted into the BED (Browser Extensible Data) format with the bedtools bamtobed tool [Quinlan and Hall 2021], and then modified for QuantifyPoly(A) input BED type files with the homemade R script. Volcano plots were generated in R. Venn diagrams were generated in R using the VennDiagram” package. Gene Ontology (GO) analysis was performed using “AnnotationDbi”, “org.At.tair.db”, “biomaRT”, “clusterProfiler” packages in R.

Gene expression level was performed on the sum of all filtered PAT-seq reads of the gene (QuantifyPolyA count option set to 10). DE genes were defined based on total transcript levels using the DESeq2 package (version 1.36.0) with the fold change level set > 2 and discovery false rate set to 0.05.

### Visualization of polyadenylation sites

For the visualization of cleavage and polyadenylation sites, modified BAM files were prepared. First, all three separate BAM files from a single genotype (from the Hisat2 alignment), were merged together into one file with the samtools merge tool [Li *et al*. 2009]. Then, merged BAM files were sorted with the samtools sort tool, and converted into the BED format with the bedtools bamtobed tool [Quinlan and Hall 2021]. Next, original alignment coordinates were exchanged for the coordinates of the first nucleotide downstream of alignment (this is the position of the first “A” in the poli(A) tail), and separated into two strands specific “plus” and “minus” files, with the homemade R script. Modified BED files were then converted back into the BAM files with the bedtools bedtobam tool [Quinlan and Hall 2021]. For each BAM file an index file was prepared with the samtools index tool [Li *et al*. 2009]. For the final visualization, BAM files were loaded to the IGV genomic browser [Robinson *et al*. 2011].

## RESULTS

### *AtCFI59* and *AtCFI68* are expressed ubiquitously in plant organs

In order to characterize the function of the CFI complex in plants, all the genes encoding homologues of the human CFIm subunits were identified in the *A. thaliana* genome. In Arabidopsis there are two genes, *AtCFI25a* and *AtCFI25b*, encoding a homolog of human CFIm25, and single genes, *AtCFI59* and *AtCFI68*, encoding homologs of human CFIm59 and CFIm68, respectively (Supplementary Fig. S1, S2). These genes were suggested earlier to encode proteins similar to human CFIm subunits [Hunt *et al*. 2012]. We previously characterized the Arabidopsis AtCFI25a protein [Zhang *et al*. 2022]. Here we focused on the two other potential subunits of the plant CFI complex. First, transcript accumulation of *AtCFI59* and *AtCFI68* was analysed in different plant organs, *i.e.,* cauline leaves, rosette leaves, roots, inflorescences, and seedlings. Primers corresponding to the unique sequences of the two genes (as shown in Fig. 1A, Supplementary Table S1) were designed. Transcripts of *AtCFI59* and *AtCFI68* were detected in all analysed organs of adult Arabidopsis plants, as well as in seedlings (Fig. 1A). Next, to investigate promoter activity of the *AtCFI59* and *AtCFI68*, intergenic regions upstream of the start codons of *AtCFI59* and *AtCFI68* were amplified and fused with the GUS marker gene, *AtCFI59_prom_*::GUS and *AtCFI68_prom_*::GUS, and transformed into wild-type plants. The activity of *AtCFI59* and *AtCFI68* promoters, was detected in all Arabidopsis organs analysed. Comparatively higher activity of the promoters was observed in anthers and pistils as well as in meristematic regions of the seedling (Fig. 1B), suggesting a special role of Arabidopsis CFI activity in reproductive organs and cell proliferation. Taken together, the mRNA expression profiles for both genes were similar. It is worth noting that the promoter activity of *AtCFI25a* spatiotemporally overlapped that of *AtCFI59* and *AtCFI68,* in reproductive organs and meristematic regions [Zhang *et al*. 2022].

**Figure 1.**
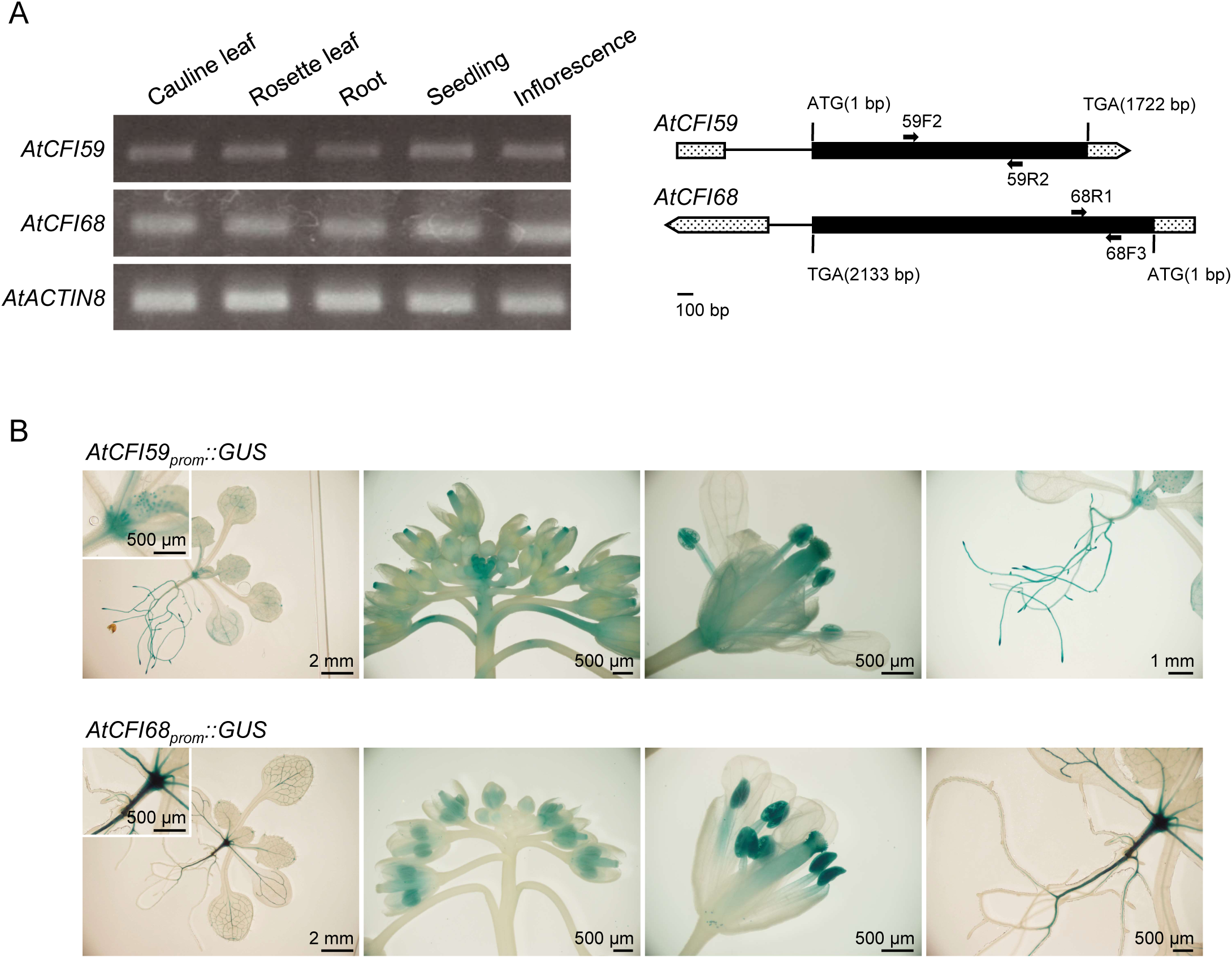
Transcripts and promoter activity of *AtCFI59* and *AtCFI68* were detected in various organs of Arabidopsis. (A) Semi-quantitative RT-PCR analyses of *AtCFI59* and *AtCFI68* transcripts in different organs of Arabidopsis (left panel) and a schematic diagram of *AtCFI59* and *AtCFI68* transcripts (right panel). Total RNA for PCR analyses was collected from subjected samples: cauline leaf, rosette leaf, root, and inflorescence were collected at 24 DAS, seedling was collected at 7 DAS. *AtACTIN8* was used as a control. Primer sequences are described in Supplementary Table S1. For the schematic diagram, exons and introns are shown in black boxes and black lines, respectively. 5’ UTR and 3’ UTR are shown in gray boxes and gray arrowed boxes, respectively. Black arrows illustrate approximate positions of primers used for RT-PCR in (A). (B) Promoter activity of *AtCFI59* and *AtCFI68* was detected in meristematic tissues and floral organs. Representative image of Arabidopsis aerial organs and root, from plants transformed with constructs expressing reporter genes, *AtCFI59_prom_:: GUS* (upper panels) and *AtCFI68_prom_::GUS* (lower panels). Transgenic lines were confirmed to harbor single homozygous copies of the reporter transgene. Note that plants expressing *AtCFI59_prom_::GUS* displayed strong promoter activity of *AtCFI59* in meristematic domains of the apical meristem, stigma, anther, root and root tip. Plants expressing *AtCFI68_prom_::GUS* displayed strong promoter activity of *AtCFI68* in meristematic domains of the apical meristem, leaf veins, anther, pistil, pollen, and primary root. Subjected samples: seedling and root were collected at 14 DAS, inflorescence and flower were collected at 24 DAS. Lengths of scale bars are noted in the figure.

### Null alleles of single *AtCFI59* and *AtCFI68* mutants display a wild-type phenotype

To further characterize the biological functions of *AtCFI59*, two T-DNA insertion lines for *AtCFI59* (SALK_090357, SALK_128104, hereafter called *atcfi59-1* and *atcfi59-2*, respectively) were identified to establish homozygous mutants. The exact insertion sites of T-DNAs are shown schematically in Supplementary Fig. S3A. RT-PCR analysis showed detectable transcript accumulation of *AtCFI59* gene in *atcfi59-2* but not in *atcfi59-1* (Supplementary Fig. S3C). For further analyses, the *atcfi59-1* allele was used as a null allele. As for *AtCFI68*, three T-DNA insertion lines (GABI_217C05, GABI_213F12, and SALK_036546, hereafter called *atcfi68-1*, *atcfi68-2,* and *atcfi68-3*, respectively) were identified to establish homozygous insertion mutants. The exact insertion sites of T-DNAs are shown schematically in Supplementary Fig. S3A. RT-PCR analysis showed transcript accumulation for *AtCFI68* in *atcfi68-1* and *atcfi68-2*, but not in *atcfi68-3* (Supplementary Fig. S3C). Therefore, *atcfi68-3* was used as a null allele.

The aerial structure of the single *atcfi59-1* and *atcfi68-3* mutants displayed morphology similar to that of wild-type plants (Fig. 2A, B), while *atcfi59-1* displayed slight defect in root morphology (Supplementary Fig. S4B). The flowering time of *atcfi59-1* and *atcfi68-3* was also similar to wild type (Supplementary Fig. S4B). Although pollen viability and fertility were normal for both *atcfi59-1* and *atcfi68-3* (Supplementary Fig. S4A), detailed analyses revealed slightly shorter siliques for *atcfi59-1* (Fig. 2C). The altered root phenotype only observed in *atcfi59* but not in *atcfi68* might be related to the different pattern observed in the promoter activity.

**Figure 2.**
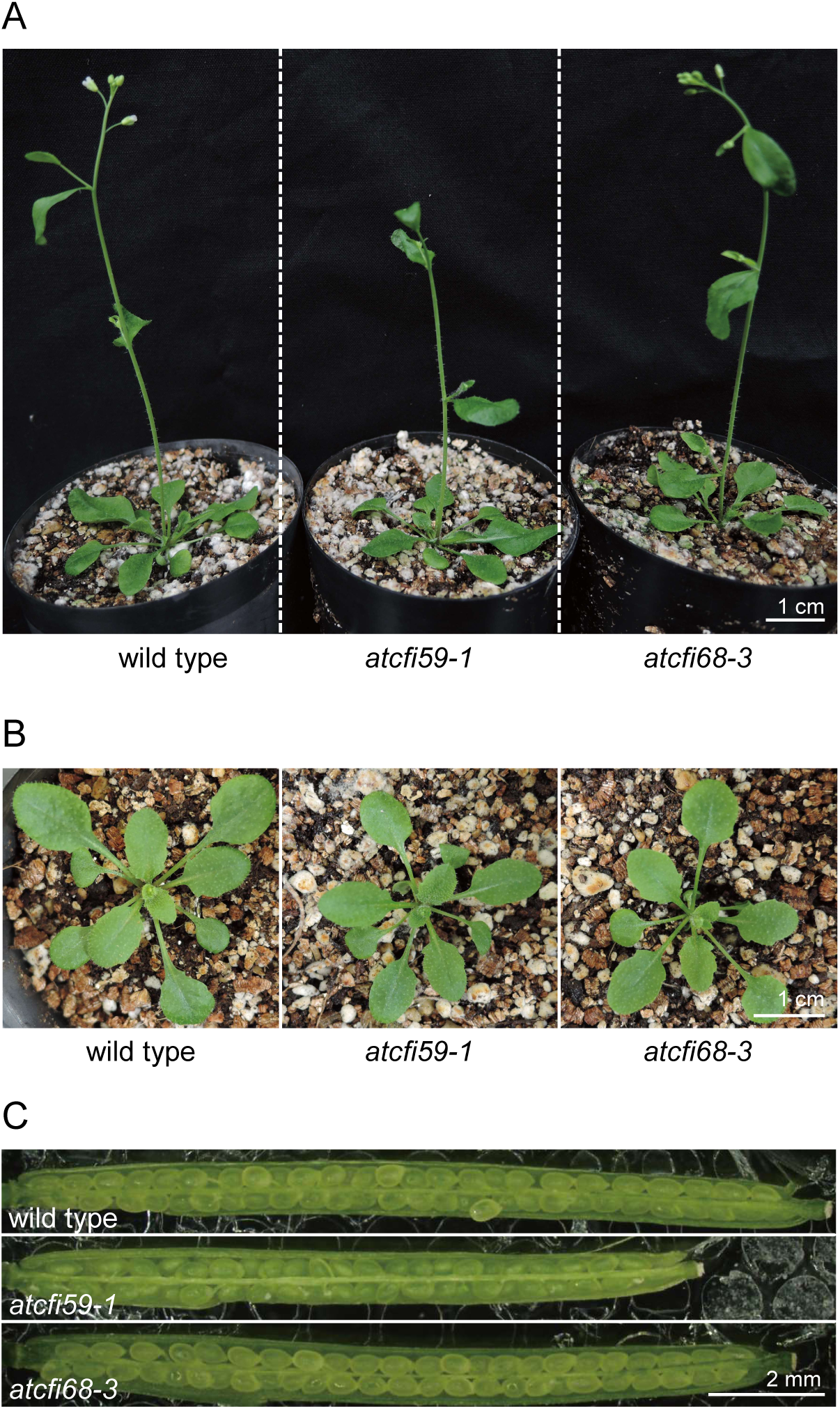
The loss of function alleles *atcfi59-1* and *atcfi68-3* display no obvious morphological differences compared to wild type. The phyllotaxy, number of organs, pigmentation, fertility of the two alleles were similar to those of the wild type. (A) Aerial structure of wild type, *atcfi59-1*, and *atcfi68-3*. (B) Rosette morphology and phyllotaxy of wild type, *atcfi59-1*, and *atcfi68-3*. Note that *atcfi59-1* and *atcfi68-3* displayed no detectable deformities. (C) *atcfi59-1* and *atcfi68-3* displayed no differences in seed formation and fertility. *atcfi59-1* showed slightly shorter siliques. Lengths of scale bars are noted in the figure.

### Loss of CFI function phenotype of *atcfi59 atcfi68* double mutant plant

Next, *atcfi59-1* and *atcfi68-3* were crossed to obtain plants harboring homozygous loss of function mutations for both *AtCFI59* and *AtCFI68* genes. The double null *atcfi59-1 atcfi68-3* mutant, displayed a severe phenotype, displaying a smaller aerial structure and shorter siliques with a smaller number of seeds per silique (Fig. 3A-D). Pollen of the double null mutant was viable, yet the anthers were smaller and contained less pollen grains (Fig. 3E). Detailed comparison of flower development from flower stage 12 to 15 [Smyth *et al*. 1990] revealed that *atcfi59-1 atcfi68-3* double mutant produced smaller flowers with undeveloped petals, shorter stamens, and elongated stigmatic papillary cells (Fig. 4). Notably, this phenotype was identical to the previously characterized *atcfi25a-3* mutant [Zhang *et al*. 2022].

**Figure 3.**
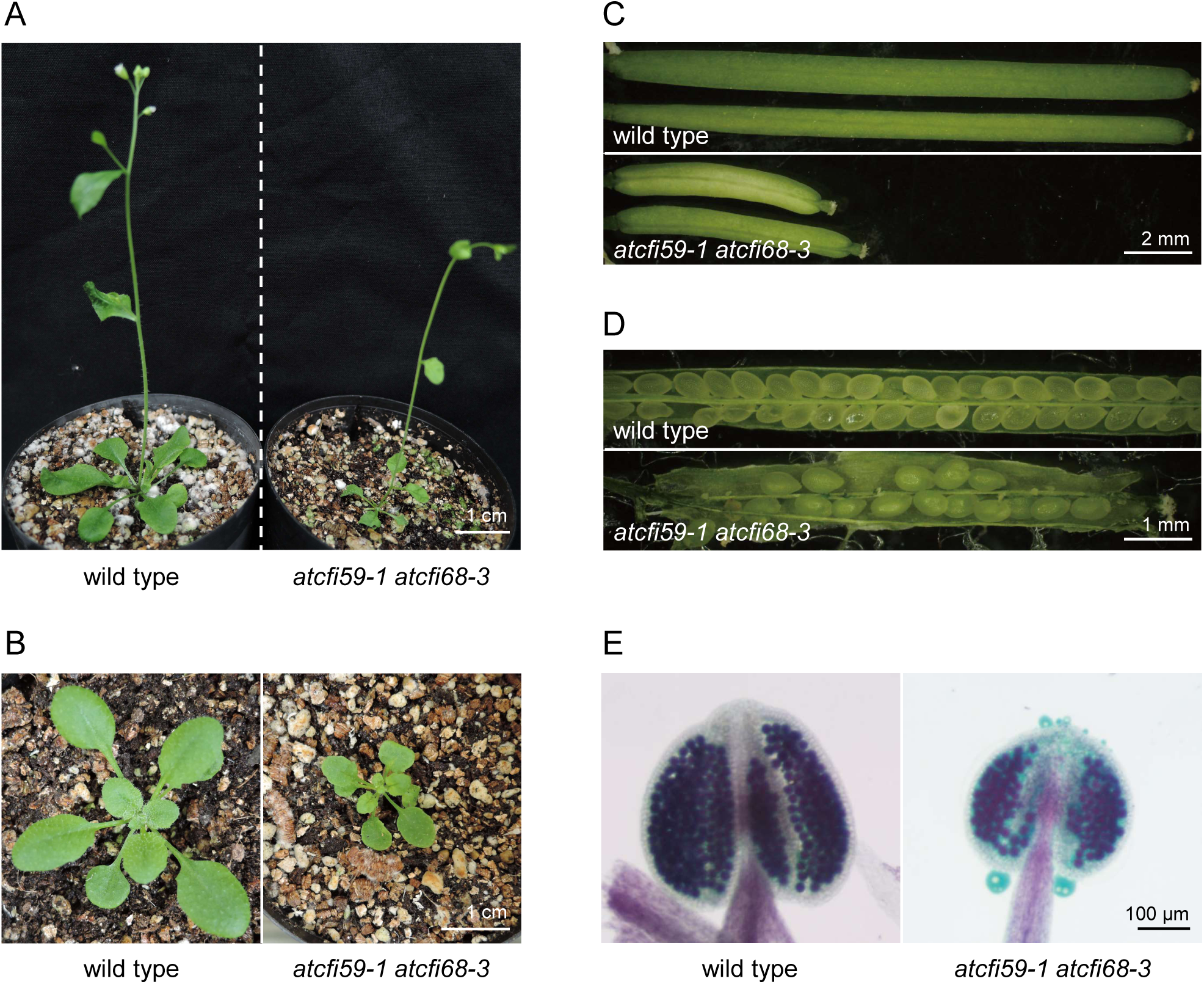
*atcfi59 atcfi68* double mutant displayed pleiotropic defects. (A) Aerial structure of wild type and *atcfi59-1 atcfi68-3*. Double mutant was confirmed to carry homozygous null alleles for *atcfi59-1* and *atcfi68-3*. (B) Dwarf phenotype of *atcfi59-1 atcfi68-3.* The phyllotaxy of *atcfi59-1 atcfi68-3* was similar to wild type. Note that *atcfi59-1 atcfi68-3* displayed pleiotropic defects identical to those observed in *atcfi25a-3* [Zhang *et al*. 2022]. Subjected samples: 25 DAS (A) and 21 DAS (B). (C) *atcfi59-1 atcfi68-3* displays short siliques. (D) *atcfi59-1 atcfi68-3* displayed low fertility and less seeds, likely due to the short stamen. (E) Alexander’s staining showed no obvious difference in the viability between the pollen of wild type and *atcfi59-1 atcfi68-3*. Note that *atcfi59-1 atcfi68-3* had smaller anther with less pollen. Lengths of scale bars are noted in the figure.

**Figure 4.**
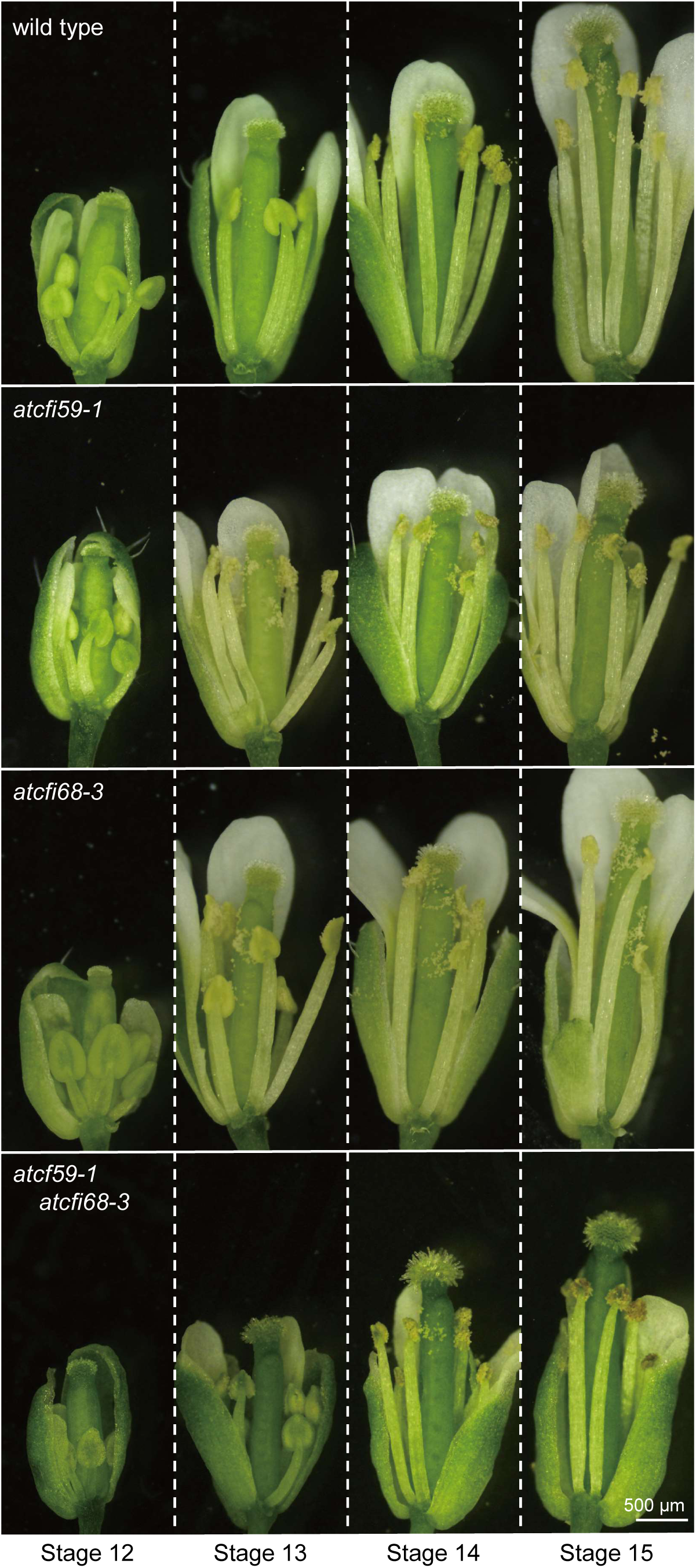
Floral morphology of wild type, *atcfi59-1*, *atcfi68-3*, and *atcfi59-1 atcfi68-3* double mutants. Representative flowers corresponding to floral stages 12 to 15 from left to right are shown [Smyth *et al*. 1990]. *atcfi59-1* showed slightly smaller flower in stage 14, 15, while *atcfi68-3* showed similar flower phenotype to wild type. Note that *atcfi59-1 atcfi68-3* possessed short stamens with anthers containing less pollen, and a pistil with abnormally elongated papillary cells. These morphological characteristics of *atcfi59-1 atcfi68-3* was identical to those observed for *atcfi25a-3* [Zhang *et al*. 2022]. Some petals and sepals were removed to show the flower structure. Lengths of scale bars are noted in the figure.

The fact that the phenotype of *atcfi59-1 atcfi68-3* double mutant highly resembles that of *atcfi25a-3* [Zhang *et al*. 2022] suggests that this morphological defect represents the loss of function of the AtCFI complex function. Furthermore, considering the structure of CFI complex, this eye-catching similarity suggests that AtCFI25, AtCFI59 and/or AtCFI68 co-ordinately function as a complex.

### AtCFI25a, AtCFI59, and AtCFI68 are subunits of the plant CFI

In an attempt to confirm the composition of Arabidopsis CFI, we performed reciprocal co-IP experiments using: AtCFI25a tagged with GFP at its N-terminus (GFP-AtCFI25a) expressed in the *atcfi25a-1* mutant [Zhang *et al*. 2022], AtCFI59 tagged with GFP at its N-terminus (GFP-AtCFI59) expressed in the *atcfi59-1* mutant, and AtCFI68 tagged with GFP at its C-terminus (AtCFI68-GFP) expressed in the *atcfi68-3* mutant. Proteins that co-purified with GFP-tagged bait proteins were identified by mass spectrometry (LC-MS/MS). Enriched proteins were determined by comparing proteins from GFP-tagged co-IPs against mock IPs, with anti-GFP antibodies. In all experiments, AtCFI25a, AtCFI59, and AtCFI68 were identified in addition of the tagged bait, however AtCFI25b was not detected (Fig. 5). These results and pull-down results indicate that AtCFI25a, AtCFI59, and AtCFI68, likely compose the CFI complex in Arabidopsis (Fig. 5, Supplementary Fig. S5).

**Figure 5.**
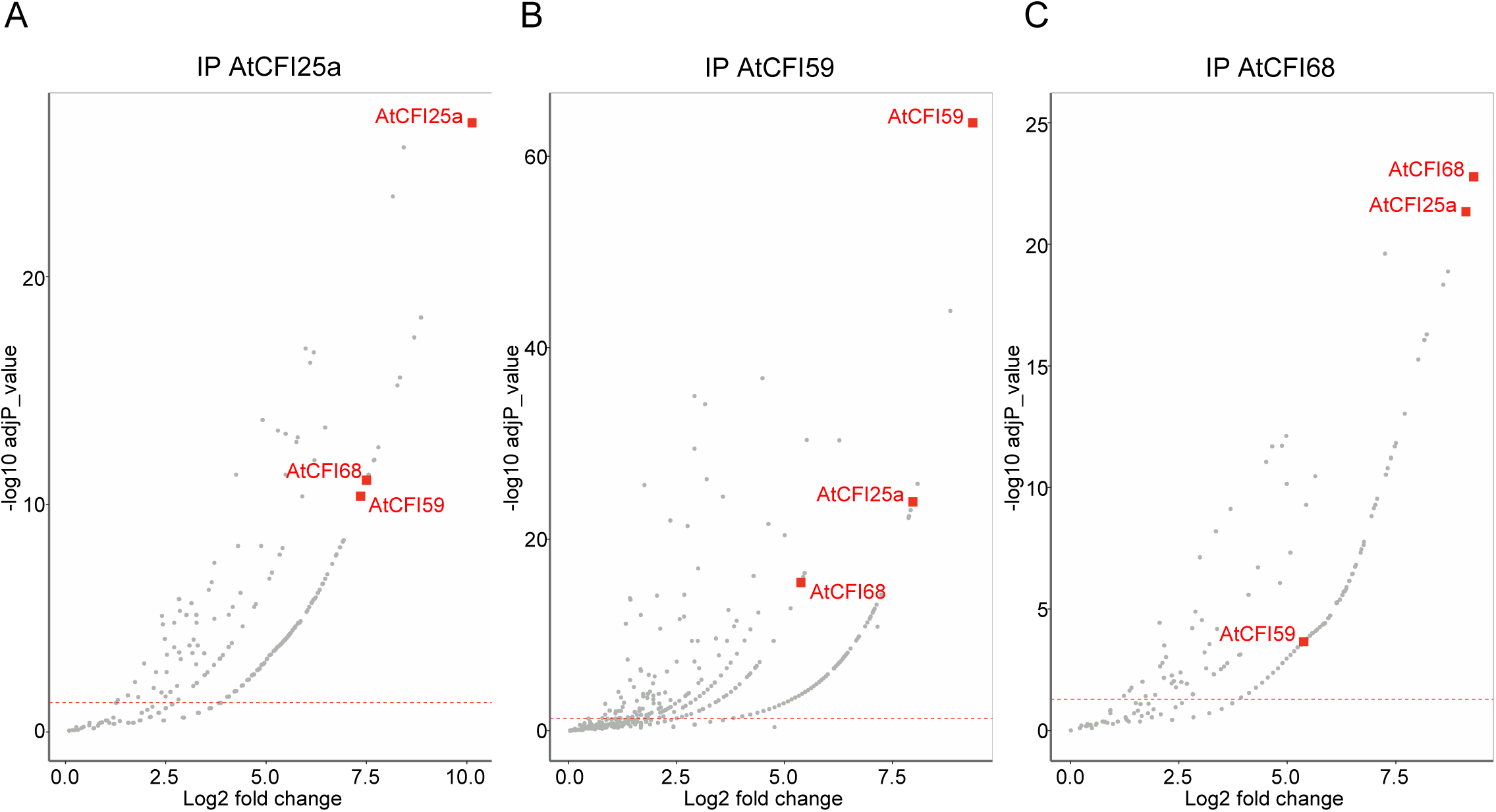
Identification of Arabidopsis CFI subunits by reciprocal co-immunoprecipitation mass-spectrometry analyses. Volcano plots showing protein that co-purified with (A) GFP-AtCFI25a, (B) GFP-AtCFI59, and (C) AtCFI68-GFP. The bait proteins are described at the top of each plot. The tagged bait protein was expressed in wild-type plants to generate extract to be subjected for the analyses. Wild type extract and anti-GFP antibody was used as control extracts. Significantly enriched proteins are above the dotted red lines (adjp < 0.05).

In order to comprehend the architecture of plant CFI complexes, we next set out to analyse the interactions between the three subunits Förster resonance energy transfer (FRET) - fluorescence lifetime imaging microscopy (FLIM). FLIM-FRET efficiency was calculated to quantify the direct protein interactions by measuring the decrease of the fluorescence lifetime of a donor protein in the presence of an acceptor protein, *i.e.*, donor: AtCFI59-GFP (Fig. 6A, B) or AtCFI68-GFP (Fig. 6C, D); acceptor: RFP-AtCFI25a, AtCFI68-RFP, and AtCFI59-RFP (Fig. 6A, B) or RFP-AtCFI25a, AtCFI59-RFP, and AtCFI68-RFP (Fig. 6C, D). RFP-PAPS1 (Arabidopsis POLY(A) POLYMERASE 1) was used as negative acceptor protein control. The fluorescence lifetime of AtCFI59-GFP and AtCFI68-GFP was reduced when RFP-AtCFI25a was co-expressed, but not when RFP-PAPS1 was co-expressed (Fig. 6B, D). These results show that AtCFI59 and AtCFI68 are directly bound to AtCFI25.

**Figure 6.**
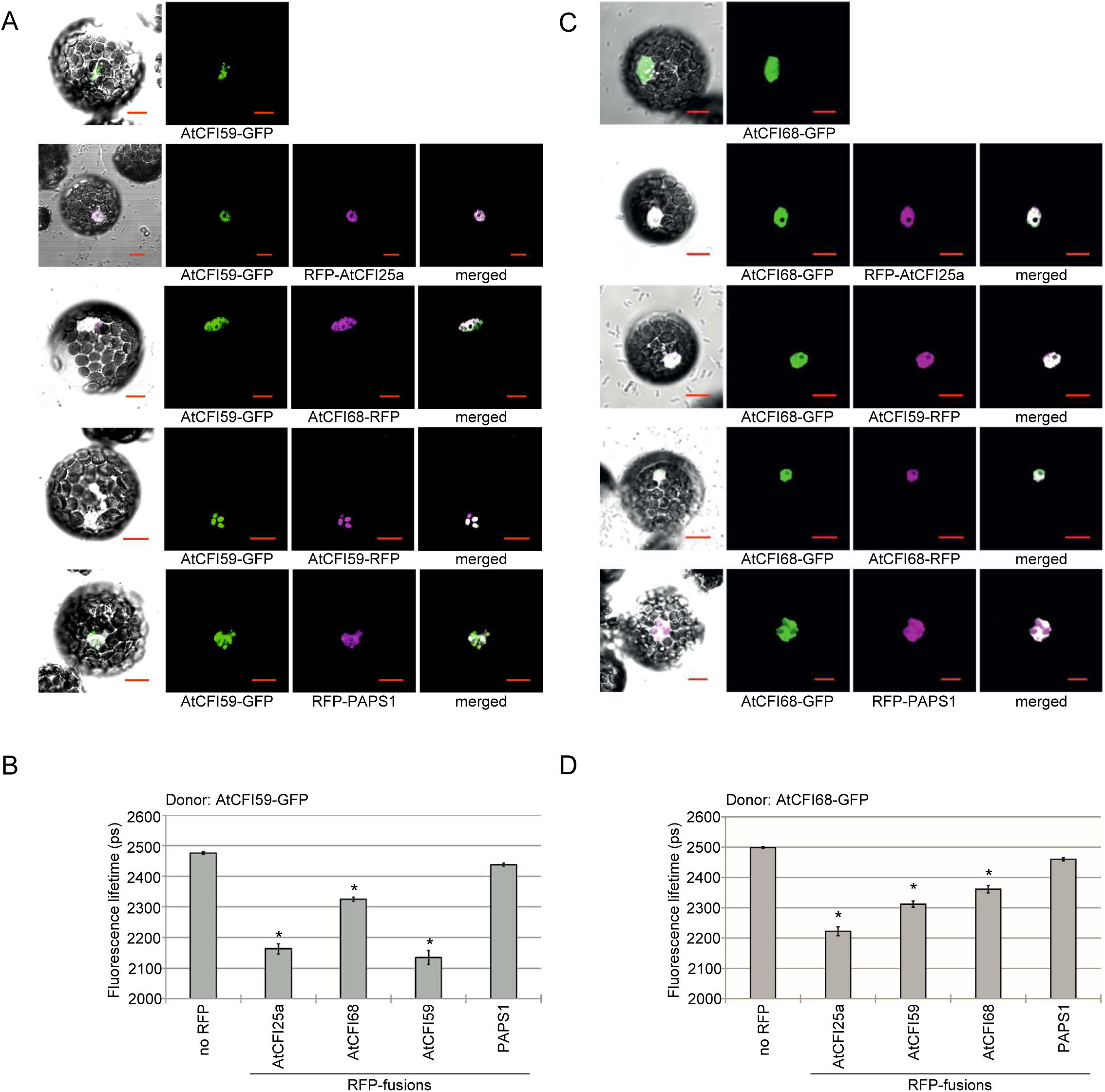
AtCFI59 and AtCFI68 interact with AtCFI25a while forming homo- or heterodimers. (A) Co-localization of AtCFI59-GFP with RFP-AtCFI25a, AtCFI68-RFP, AtCFI59-RFP, and RFP-PAPS1 in Arabidopsis protoplasts. (B) FRET-FLIM analyses in Arabidopsis protoplasts expressing AtCFI59-GFP donor protein together with RFP-AtCFI25a, AtCFI68-RFP, AtCFI59-RFP, or RFP-PAPS1 (negative control) acceptor proteins. (C) Co-localization of AtCFI68-GFP with RFP-AtCFI25a, AtCFI59-RFP, AtCFI68-RFP, and RFP-PAPS1 in Arabidopsis protoplasts. (D) FRET-FLIM analyses in Arabidopsis protoplasts expressing AtCFI68-GFP donor protein together with RFP-AtCFI25a, AtCFI59-RFP, AtCFI68-RFP, or RFP-PAPS1 (negative control) acceptor proteins. Graph presents the fluorescence lifetime of the donor protein in picoseconds (ps). Error bars indicate the standard error of the mean, n ≥ 20. Asterisks indicate significant differences (p < 0.001, Mann-Whitney-Wilcoxon test) between the control samples co-expressing the donor fused to GFP and the indicated acceptors fused to RFP to the samples expressing the donor and RFP-PAPS1 control. Lengths of scale bars equal 10 µm.

Interestingly, the fluorescence lifetime of AtCFI59-GFP and AtCFI68-GFP was also reduced upon co-expression of AtCFI59-RFP or AtCFI68 RFP (Fig. 6B, D, respectively), which strongly suggests that AtCFI59 or AtCFI68 would form heterodimers as well as homodimers.

Taken together, the results indicate that the plant CFI complex, similar to its human counterpart, is composed of three subunits: AtCFI25a, AtCFI59 and AtCFI68. The AtCFI25a subunit interacts with AtCFI59 as well as with the AtCFI68 protein. In addition, AtCFI59 and AtCFI68 were shown to interact with each other, suggesting homo- and heterodimerized forms of these proteins in the plant CFI complex.

### AtCFI59 or AtCFI68 are necessary for efficient nuclear localization of AtCFI25a

While performing FLIM-FRET experiments, we noticed that in transformed Arabidopsis protoplasts, the AtCFI59 and AtCFI68 subunits behave differently compared to AtCFI25a. GFP tagged version of both AtCFI59 and AtCFI68 were detected in the nucleus, while the GFP tagged version of AtCFI25a was detected in both the nucleus and the cytoplasm (Fig. 7A). However, when we transformed protoplasts with a mixture of AtCFI59 and AtCFI25a, or AtCFI68 and AtCFI25a, we detected a clear nuclear co-localization of the combined proteins of interest (Fig. 7B). Interestingly, AtCFI25a was predominantly detected in the nucleus, and the cytosolic signal was lost only when co-expressed with AtCFI59 or AtCFI68, suggesting that the presence of either AtCFI59 or AtCFI68 facilitates the nuclear localization of AtCFI25a.

**Figure 7.**
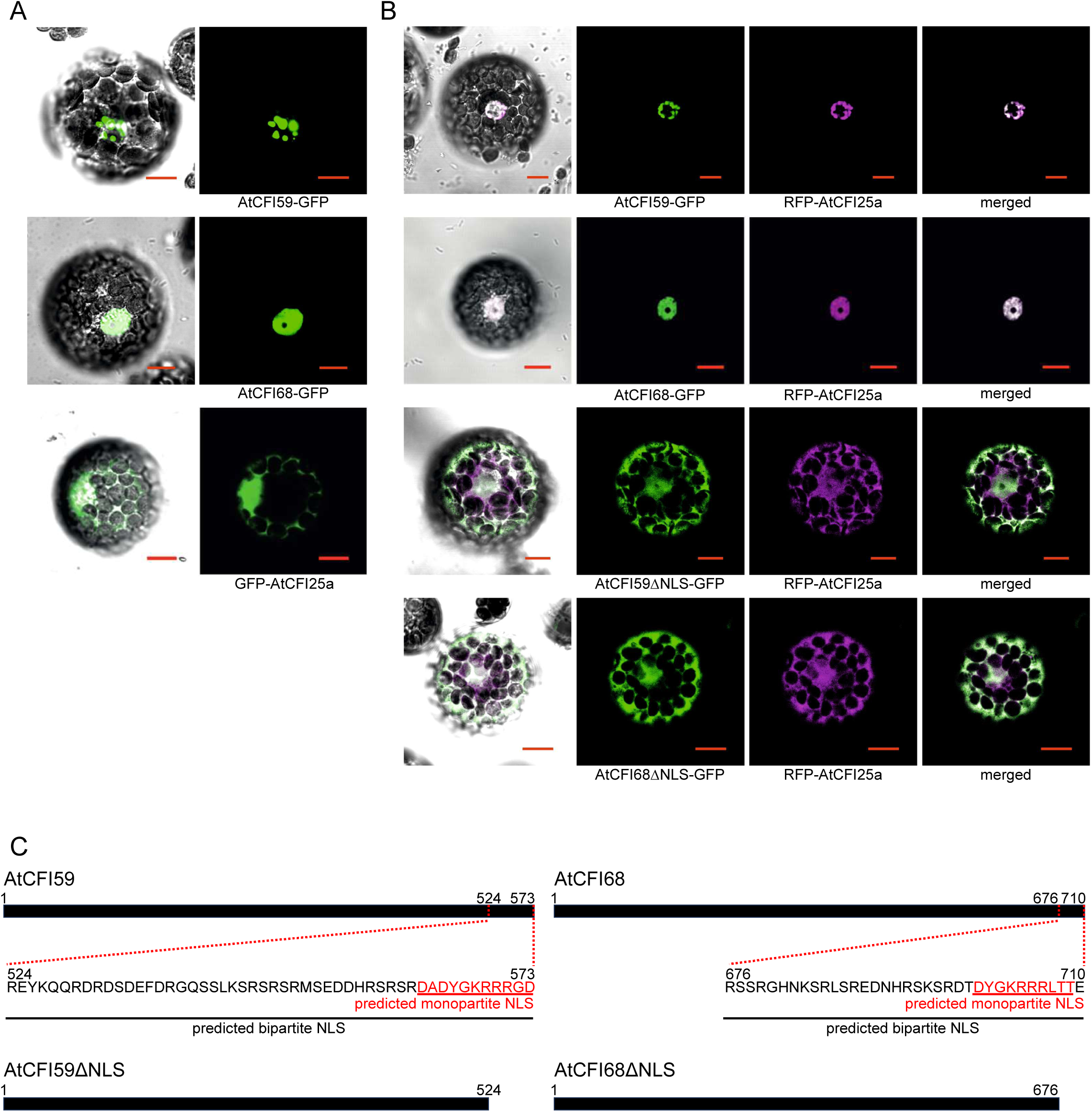
Nuclear localization of AtCFI25a requires AtCFI59 or AtCFI68. (A) Localization of GFP-AtCFI25a, AtCFI59-GFP, and AtCFI68-GFP in Arabidopsis protoplasts. AtCFI59 and AtCFI68 predominantly localize in the nucleus while AtCFI25a can be found in both nucleus and cytoplasm. (B) Localization of RFP-AtCFI25a, when co-expressed with either AtCFI59-GFP, AtCFI68-GFP, AtCFI59ΔNLS-GFP, or AtCFI68ΔNLS-GFP. Note that cytoplasmic signal of RFP-AtCFI25a were lost in the upper two rows while it increased in the lower two rows. The presence of either AtCFI59 or AtCFI68 in the nucleus facilitates nuclear localization of AtCFI25a. (C) Schematic diagram of AtCFI59 and AtCFI68 with predicted NLS amino acid sequence and position in numbers. NLS deleted protein are shown below. Red letters and line show predicted monopartite NLS, while black show predicted bipartite NLS signals. Lengths of scale bars equal 10 µm.

We then hypothesized that the nuclear localization signals (NLS), found at the C-terminal ends of AtCFI59 and AtCFI68, might be important for the nuclear localization of these two proteins, and hence the nuclear localization of AtCFI25a. To test this hypothesis, we analysed the NLS by utilizing the cNLS Mapper [Kosugi *et al*. 2009] and predicted a bipartite NLS within the region of the last forty-nine amino acid residues for AtCFI59, as well as the last thirty-four amino acid residues for AtCFI68. Constructs lacking the predicted bipartite NLS in AtCFI59 and in AtCFI68 (AtCFI59ΔNLS and AtCFI68ΔNLS, respectively, Fig. 7C, lower panels) were generated and co-transformed with AtCFI25a into the protoplasts for localization studies. AtCFI59ΔNLS and AtCFI68ΔNLS localized mostly to the cytoplasm, with detectable diffused signals in the nucleus. The merged images with the localization of AtCFI25a indicated that AtCFI59 or AtCFI68 lacking the predicted NLS will no longer facilitate the predominant nuclear localization of AtCFI25a (Fig. 7B).

These results show that Arabidopsis CFI59 and CFI68 both use their NLS located at their C-terminal ends to be localized to the nucleus, through active transportation. In contrast, the Arabidopsis CFI25 is stably localized to the nucleus through interacting with CFI59 and/or CFI68.

### AtCFI determines the 3′ UTR length of various genes

To characterize the function of plant CFI complex, from the viewpoint of its regulation on global polyadenylation events, poly(A) tag sequencing (PAT-seq), a high-throughput method for global determinations of poly(A) site choice, was applied in Arabidopsis [Pati *et al*. 2015]. The variation of polyadenylation site selection in selected genetic backgrounds, *i.e., atcfi25a*, *atcfi25b*, *atcfi25a atcfi 25b* double mutant, *atcfi59*, *atcfi68*, *atcfi59 atcfi68* double mutant, relative to wild type was analysed.

The sequencing results were processed using QuantifyPoly(A) [Ye at al. 2021], a complete and straightforward pipeline for quantifying poly(A) site profiles from high-throughput sequencing data. The clustered polyadenylation sites were then analysed for changes in frequency of occurrence in individual mutants relative to control plants. An increase in polyadenylation frequency in the direction of the transcription start site was identified as increased selection of proximal polyadenylation sites, and an increase in polyadenylation frequency in the opposite direction was identified as increased selection of distal polyadenylation sites.

Widespread changes in the choice of the polyadenylation site were observed. More than 4000 genes showed a shift to the proximal PA site in *atcfi25a* (Fig. 8A), and the *atcif25a atcif25b* double mutant (Fig. 8C), resulting in shortened 3’ UTRs, whereas around 400 transcripts showed a shift to the distal CPA site, resulting in lengthened 3’ UTRs. Overlapping the genes with altered CPA site in both mutants revealed a high degree of overlap (Supplementary Fig. S6A). In line with this, the *atcfi25b* mutant showed almost no changes in the CPA site choice (Fig. 8B). In the *atcfi59* and *atcfi68* single mutants, a limited number of transcripts showed alterations in the choice of the CPA site (Fig. 8 D, E) whereas almost 5000 genes showed a change in CPA site choice in the double mutant (Fig. 8 F, Supplementary Fig. S6C). Again, a predominant shift to the proximal CPA site was observed. Notably, comparison of the genes with aberrant polyadenylation in the *atcfi25a* mutant, the *atcfi25a atcfi25b* double mutant, and the *atcfi59 atcfi68* double mutant revealed an almost complete overlap (Supplementary Fig. S6B). Notably, the extent of changes in CPA site choice and thus 3’ UTR length in the mutants correlated with the severity of morphological defects in the mutants.

**Figure 8.**
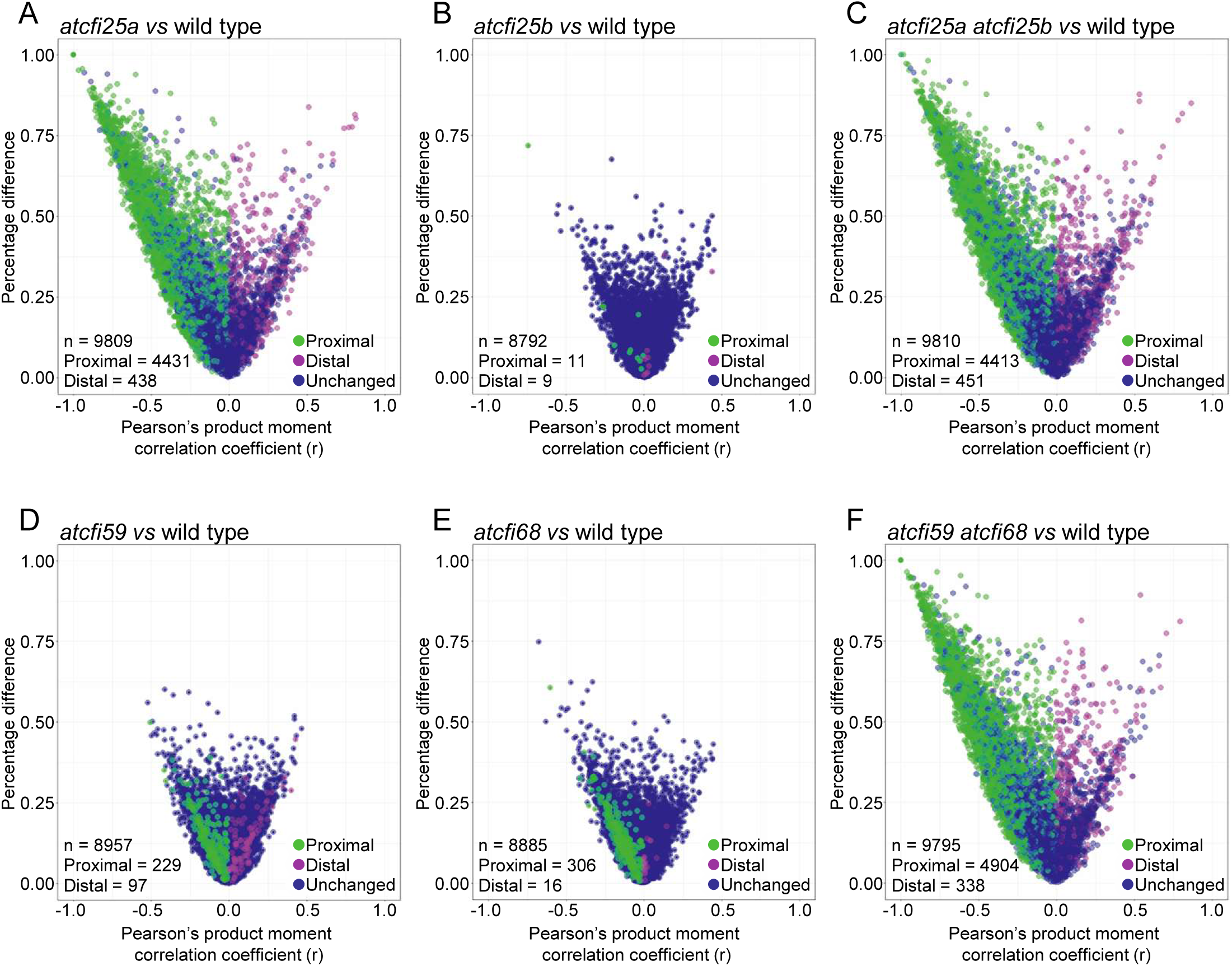
Differential usage of CPA sites in *atcfi25a*, *atcfi25b*, *atcfi59*, *atcfi68*, *atcfi25a atcfi25b* double mutant, and *atcfi59 atcfi68* double mutant in comparison to wild type. Scatter plots of the Pearson’s product moment correlation coefficient (r) and percentage difference of poly(A) site clusters (PACs) were determined by QuantifyPoly(A) program [Ye *et al*. 2021]. Green and magenta dots represent PAC usages showing significant difference between genotypes, in favour for proximal and distal distribution, respectively. Dark purple dots represent those showing no significant difference between genotypes. n: total number of identified PACs in a given comparison.

Taken together, despite the high sequence resemblance between *AtCFI25a* and *AtCFI25b* [Zhang *et al*. 2022], AtCFI25b is likely not the component of the Arabidopsis CFI complex, under our growth condition. AtCFI59 and AtCFI68 seems to share a redundant, or partially redundant, function in maintaining the overall 3’ UTR length. Since the changes observed in the 3’ UTR length of global genes are similar (more than 80%) among *atcfi25a*, *atcfi25a atcfi25b* double mutant, and *atcfi59 atcfi68* double mutant, this alteration in 3’ UTR length would be due to the loss of CFI complex function in Arabidopsis (Supplementary Fig. S6).

### CFI regulates polyadenylation site selection of its own subunits

Among the genes showing changes in CPA site choice in the mutants were the genes encoding the CFI subunits themselves. Although the 3’ UTR length pattern of *AtCFI25a*, *AtCFI59*, *AtCFI68* transcripts were similar in wild type, *atcfi25b*, *atcfi59*, and *atcfi68* mutants, the pattern was greatly altered in *atcfi25a*, *atcfi25a atcfi25b* double mutant, and *atcfi59 atcfi68* double mutant (Fig. 9). The absence of neither *AtCFI59* nor *AtCFI68* function alone did not have any effect on CPA site regulation of *CFI* subunit transcripts (Fig. 9A, B). However, simultaneous loss of *AtCFI59* and *AtCFI68* function resulted in a reduction of the number of CPA sites, and consequently shortening of 3’ UTR length, identical to what was observed in loss of *AtCFI25a* function, or simultaneous loss of *AtCFI25a* and *AtCFI25b* function (Fig. 9A-E). This was in line with the semi-quantitative RT-PCR analyses performed on the 3’ UTR regions of *AtCFI25a*, *AtCFI25b*, *AtCFI59*, and *AtCFI68* transcripts (Supplementary Fig. S7). These results support the previous demonstration of a self-regulatory role of *AtCFI25a* function in Arabidopsis CFI subunit transcription [Zhang *et al*. 2022].

**Figure 9.**
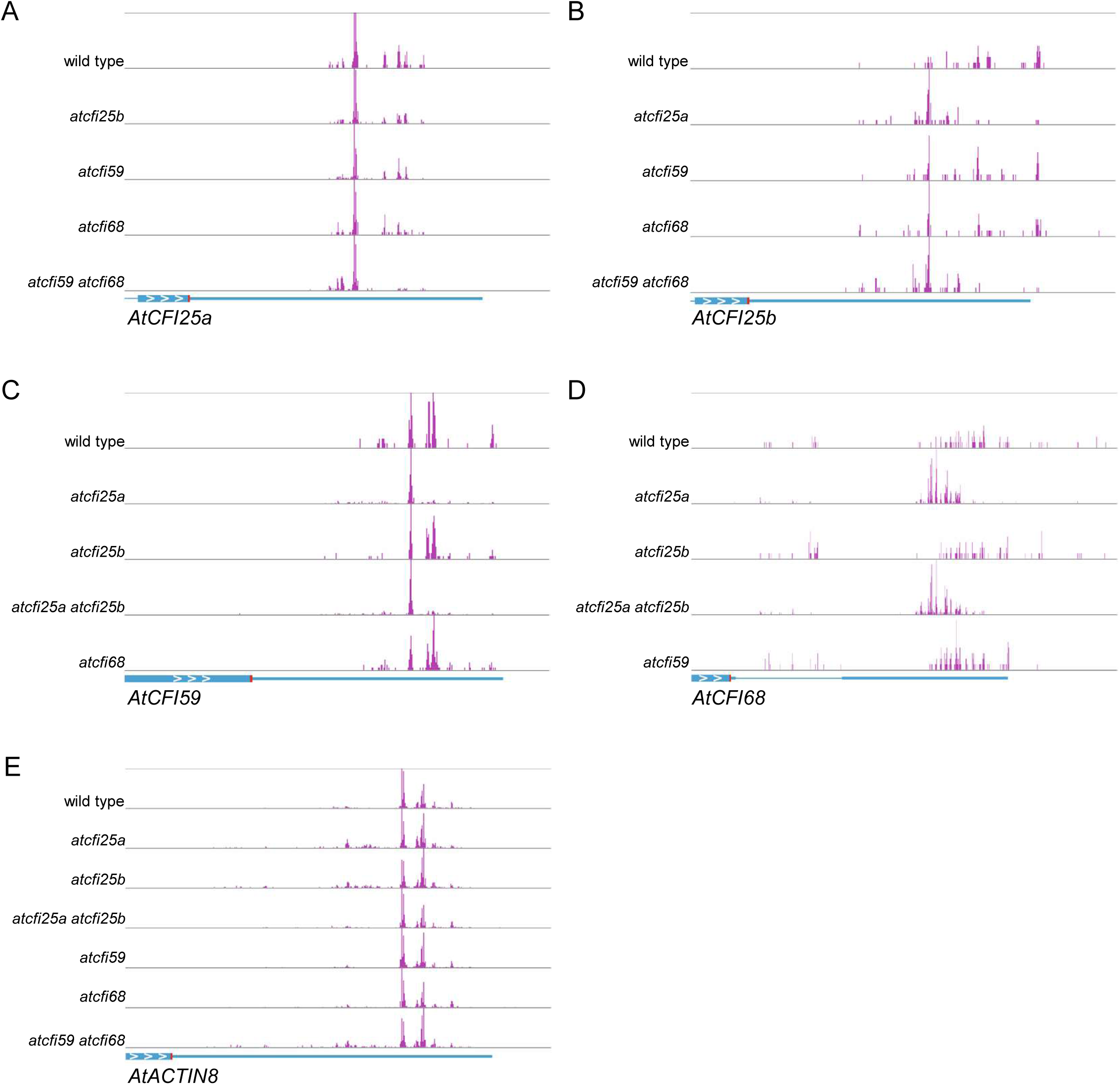
CFI function is essential to maintain diverse cleavage and polyadenylation site selection pattern for CFI subunit encoding genes. PAT-seq analyses conducted for transcripts of (A) *AtCFI25a*, (B) *AtCFI25b*, (C) *AtCFI59*, (D) *AtCFI68*, and (E) *AtACTIN8*, reveals a self-regulation function of CFI on its subunit encoding genes. A diagram of the analysed transcript is shown at the bottom of each panel. The genotype of the analysed samples is noted on the left side of the panel. Each peak shown in magenta represents the position and frequency of the cleavage and polyadenylation site derived from PAT-seq analyses. Note that loss of CFI function mutants that show severe pleiotropic deformity (such as *atcfi25a*, *atcfi25a atcfi25b* double mutant, and *atcfi59 atcfi68* double mutant) correspond to those samples that give rise to obvious pattern changes in cleavage and polyadenylation site selection (also see Supplementary Figure S7). The extent of self-regulation differs among genes. Whole or part of the last exon is shown as blue box with arrowheads, intron is shown as thin blue line, 3’ end of annotated known genomic structure is shown as thick blue line. Stop codon is shown as red box. White arrowheads indicate the direction of transcription.

In summary, plant CFI complex was shown to maintain the 3’ UTR length distribution of global genes, its role in CPA site regulation (Fig. 8, Fig. 9). Furthermore, AtCFI59 and AtCFI68, were shown to be involved in a self-regulatory role of CFI subunit transcription in plants, in a manner similar to AtCFI25a [Zhang *et al*. 2022].

## DISCUSSION

### Plant CFI composition and its heterogeneity

CFIm25 has been reported as the essential subunit for CFIm formation in mammals [Yang *et al*. 2011], forming homodimers facilitating the binding interface to the pre-mRNA at FUEs [Venkataraman *et al*. 2005]. Previously we have reported that the plant AtCFI25 would possess all characters required to function in the CFI subunit counterpart in Arabidopsis [Zhang *et al*. 2022]. In analogy to CFIm25 of the homodimer would interact with either CFIm59 or CFIm68 in mammals to form a tetramer, here we demonstrated that AtCFI25a, AtCFI59, AtCFI68 are the three components that interact with each other in plants. This is in line with our *in silco* modelling suggesting that the Arabidopsis CFI complex may exhibit similar feature, where AtCFI25a is forming a dimer and interacting with either AtCFI59 or AtCFI68 [Zhang *et al*. 2022]. Interestingly, AtCFI25b, which is the second putative Arabidopsis CFI25 protein, is not a subunit of the AtCFI complex, suggesting that it is not involved in regulation of polyadenylation of plant RNAPII transcripts, at least under the conditions and developmental stages analyzed in this study.

In this paper we provide experimental data supporting the idea that architecture of the Arabidopsis CFI complex is similar to those described in mammals. First, co-IP always detected all of the three subunits concentrated in the purified elute, when either AtCFI25a, AtCFI59, or AtCFI68 was used as a bait, (Fig. 5). AtCFI25a was also shown to bind AtCFI59 and AtCFI68 independently on one-to-one binding assays (Supplementary Fig. S5) [Zhang *et al*. 2022].

Second, fluorescence confocal microscopy combined with transient expression of AtCFI subunits in plant protoplasts showed that AtCFI25a co-localized with AtCFI59 or AtCFI68, individually, in the nucleus (Fig. 6A, C). FLIM-FRET experiments further clarified the interaction between AtCFI25a with AtCFI59, and AtCFI25a with AtCFI68. Interestingly, AtCFI59 and AtCFI68 subunits were also shown to interact directly with each other to form homo- and heterodimers (Fig. 6B, D), which has not been reported among mammalian CFI subunits. These are very interesting observations that can have functional implications. In mammals, one of the working models of CFI complex action proposes that CFIm can loop the intervening sequence between two UGUA elements through the dimerization of CFIm25, affecting the choice of the CPA sites [Yang *et al*. 2011]. Direct interaction between two larger subunits of the CFI complex in plants adds another layer of diversity to this model, creating a possibility of interaction between multiple variations of CFI complexes on one pre-mRNA molecule. Taken together, this can lead to looping out different parts on one pre-mRNA molecule that expands the regulation of recognition of polyadenylation signals along a given pre-mRNA.

Third, two other Arabidopsis polyadenylation factors, AtFIP1 and AtCPSF30, were concentrated in all the co-IP elute (Supplementary Table S4). AtFIP1 and AtCPSF30 are both elements of the plant CPSF complex. It was shown that the N-terminal 137 amino acids of the 1196 amino acid AtFip1, also known as AtFip1(V), directly interacts with AtCPSF30 and AtCFI25a [Addepalli and Hunt, 2007]. This raises a question how AtFIP1 would interact with the CFI complex while AtCFI25a is already bound to AtCFI59 and/or AtCFI68. On the other hand, in mammals, FIP1 directly interacts with CFIm68 [Zhu *et al*. 2018] and this interaction is suggested to activate polyadenylation factors, while CFIm59 in cooperation with U1 snRNP can suppress premature CPA in a process known as telescripting [So *et al*. 2019].

Including models where AtFIP1 would replace AtCFI59 and/or AtCFI68 in the AtCFI, which could act as a molecular switch to change the mode of CFI function, there is much to be studied to understand the CPA regulation at the 3’ ends of mRNA, despite the general similarity between plant and animal CFI complexes.

### Plant CFI localization and formation

The assembly and formation of CFI complex has been long obscure. In order to shed light to this question, the localization of Arabidopsis CFI subunits in combinations with and without deletions in the NLS of AtCFI59 and AtCFI68, was characterized in detail.

The confocal microscopy images in the protoplasts demonstrated that plant AtCFI25a was predominantly localized to the nucleus only when AtCFI59 or AtCFI68 was co-expressed (Fig. 7A), suggesting that AtCFI59 and AtCFI68 facilitates the nuclear localization of AtCFI25a. The deletion of NLS located at the C-terminal ends of AtCFI59 and AtCFI68 (AtCFI59ΔNLS and AtCFI68ΔNLS, respectively) demonstrated that either protein was no longer able to localize to the nucleus on its own. When AtCFI59ΔNLS and AtCFI68ΔNLS was expressed in combination with AtCFI25a, the smallest subunit of the CFI complex also failed to localize to the nucleus.

These results imply that the CFI complex would be formed in the cytoplasm and transported to the nucleus by utilizing the NLS of AtCFI59 and/or AtCFI68. This hypothesis would allow other ideas to be considered, such as a mechanism to serve as a quality control checkpoint for the CFI complex. assembly to for instance regulate the stoichiometry of its assembly, necessary for the proper function of the CFI complex in CPA. Alternatively, there are other models where AtCFI59 and/or AtCFI68 localize to the nuclear in advance to stabilize or trap AtCFI25a in its vicinity, therefore carrying out CPA function (Fig. 7).

### AtCFI59 and AtCFI68 has partially redundant function in plant CFI

Loss of function analyses revealed that only the *atcfi59 atcfi68* double mutant, but neither single mutant, displayed morphological defects when compared to wild-type plant, indicating functional redundancy between *AtCFI59* and *AtCFI68*. Furthermore, the *atcfi59 atcfi68* double mutant showed pleiotropic morphological defects such as dwarf pale green aerial structure, abnormal flower structure, low fertility, which were identical to those observed earlier for *atcfi25a.* These mutants also shared a sudden loss of greenness starting in mid-stalk region, eventually expanding to the whole inflorescence. Taken together, the results suggest that these morphological defects were due to the loss of function of the Arabidopsis CFI complex.

When we looked at the molecular function of CFI subunits, PAT-seq results show that the plant CFI complex is essential to maintain the diverse 3’ UTR length distribution, seen in wild-type plants (Fig. 9). Null alleles for *AtCFI25a* or both *AtCFI59* and *AtCFI68* result in less diverse CPA patterns, favoring shorter 3’ UTR in many genes. The fact that there is a fraction of genes that favored longer 3’ UTR, which has not been reported in knock-down studies of mammalian CFI functions [Zhu *et al*., 2018], is interesting. This could be due to the unique character of 3’ UTR CPA regulation in plants, or due to the high resolution of CPA sites of the current study.

The changes in the 3’ UTR length pattern observed in *atcfi25a*, *atcfi25a atcfi25b* double mutant, and *atcfi59 atcfi68* double mutant, share high similarity as noted before (Fig. 9, Supplementary Fig. S6). The CPA sites of 45.2% genes shifted to proximal sites, while only 4.5% genes shifted to distal sites in *atcfi25a* mutants when compared to wild-type plants. In a similar manner, CPA sites of 45.0% genes shifted to proximal sites while 4.6% genes shifted to distal sites in the *atcfi25a atcfi25b* double mutant, and 50.1% genes shifted to proximal sites while 3.5% genes shifted to distal sites in the *atcfi59 atcfi68* double mutant.

In order to investigate whether there was a preference for the extent of changes, gene ontology (GO) enrichment analyses were conducted using the datasets. As shown in Supplementary Fig. S8, the genes that are involved in light regulation were among the highly affected biological processes. The processes ranged from upstream factors such as “response to light intensity”, “photosynthesis light harvesting”, and “photosynthesis light reaction”, to downstream factors physiological effects such as “stomatal movement”, “regulation of stomatal movement”, “plastid organization”, and “response to oxidative stress”. These data are in line with the phenotype we observe for these mutants. The changes could explain the less accumulation of light responding pigmentation such as anthocyanin or chlorophyll in the mutants, and the dwarfness due to altered photosynthesis. In short, the results suggest that genes responding to environmental changes highly rely on the CFI function in plants.

The mechanism to regulate the preferences of the genes that rely on CFI function in plants are yet to be discovered. The selectivity could be due to some unknown *cis* elements or a novel player that would fine tune the known regulation of CPA regulating factors at the 3’ ends. Alternatively, it could be due to variations in the component of the plant CFI complex, giving rise to a variety of CFI isoforms that could also be regulated in a spaciotemporal manner. Further experiments, however, are needed to answer these interesting questions.

### Regulation of the 3′ UTR length by AtCFI can both increase as well as decrease the accumulation of transcripts

Interestingly, the changes in length of mRNA 3’ UTRs observed in *atcfi25a*, *atcfi25a atcfi25b* double mutant, and *atcfi59 atcfi68* double mutant had bi-directional influence on the level of transcript accumulation (Supplementary Fig. S9). More than 20% of the genes with CPA site changes resulted in alteration of transcript levels, with roughly half of them causing an increase and half a decrease in transcript accumulation levels. Although in most cases the inactivation of AtCFI complex activity resulted in shortening of mRNAs at the 3’ ends, genes of the lengthened 3’ ends had both increased and decreased transcript levels, similar to those of the shortened 3’ ends. Therefore, it is clear that CFI-dependent selection of CPA sites can have different effects on accumulation of different mRNAs. Although many changes of polyadenylation sites that are detected in CFI complex mutants are not linked to transcript levels, the endogenous function of AtCFI is attributed in the machinery of transcript accumulation regulation.

Interestingly, for genes encoding the CFI subunits, proximal CPA sites were favoured upon inactivation of AtCFI complex activity, leading to shortened 3’ UTRs (Fig. 9). In consequence, the shortening of these mRNAs increased the transcript accumulation of genes (Supplementary Fig. 10). This is interesting example of self-regulation that is important for cleavage and polyadenylation of the sites controlled by the CFI complex.

### Unique functions of plant CFI

While generally mutant phenotypes (Fig. 2, Fig. 3) and the 3’ UTR length analyses on global gene transcripts (Fig. 8, Fig. 9) suggest redundant functions of *AtCFI59* and *AtCFI68*, slight differences in the rosette size (Fig. 2) or nuclear localization patterns (Fig. 6, Fig. 7) might hint that the two gene products might function differently under certain conditions. AtCFI59 is more similar to the human CFIm59 rather than AtCFI68, yet AtCFI59 and AtCFI68 show high similarity to be reported as orthologs [Hunt 2023]. It has been a quest to attribute the plant homologs to the mammalian counterparts. We therefore conducted extensive plant phylogenetic analyses to reveal that plant CFI59 and CFI68 proteins form two separate clades, harboring AtCFI59 and AtCFI68 in respective clades. Based on the similarity of AtCFI59 to the human CFIm59 we consider the respective clades to harbor respective plant counterparts of the CFI subunits (Supplementary Fig. S2).

It is interesting to note that in the case of mammals, CFIm59 and CFIm68 are reported to have different functions, such as during HIV-1 infection [Luchsinger *et al*. 2023], temporal distribution among mouse early embryogenesis [Li *et al*. 2023], and CPA patterns in *CFIm59* and *CFIm68* knockdown cell lines [Tseng *et al*. 2022]. CFIm68 has been reported to form bio-condensates in human cells induced by HIV-1 infection [Ay and Di Nunzio 2023], and furthermore, CFIm25 but no CFIm59 was found in the condensates [Luchsinger *et al*. 2023]. Whether the plant CFI complex has similar phase separation property might be worth investigating in the future.

## Supporting information

Suplementary Table S4 - Results of the co-IP experiments

## ACCESSION NUMBERS

Gene names are attributed to the existing AGI locus code in this publication as follows, AtCFI25a: At4g25550, AtCFI25b: At4g29820, AtCFI59: At1g13190, AtCFI68: At5g55670.

## ACKNOWLEDGEMENT

Authors would like to thank Dr. Agata Stepien for her help in FRET-FLIM analyses. The equipment in the Mass Spectrometry Laboratory (Institute of Biochemistry and Biophysics, Polish Academy of Science, Warsaw, Poland) was sponsored in part by the Centre for Preclinical Research and technology (CePT), a project co-sponsored by European Regional Development Fund and Innovative Economy, The National Cohesion Strategy of Poland.

## REFERENCES

1. Saini, K.S., Summerhayes, I.C. and Thomas, P. (1990) Molecular events regulating messenger RNA stability in eukaryotes. Mol. Cell. Biochem., 96, 15–23.

2. Mangus, D.A., Evans, M.C. and Jacobson, A. (2003) Poly(A)-binding proteins: multifunctional scaffolds for the post-transcriptional control of gene expression. Genome Biol., 4, 223.

3. Wigington, C.P., Williams, K.R., Meers, M.P., Bassell, G.J. and Corbett, A.H. (2014) Poly(A) RNA-binding proteins and polyadenosine RNA: new members and novel functions. Wiley Interdiscip. Rev. RNA, 5, 601–622.

4. Park, J., Kim, M., Yi, H., Baeg, K., Choi, Y., Lee, Y.S., Lim, J. and Kim, V.N. (2023). Short poly(A) tails are protected from deadenylation by the LARP1-PABP complex. Nat. Struct. Mol. Biol., 30, 330–338.

5. Apponi, L.H., Leung, S.W., Williams, K.R., Valentini, S.R., Corbett, A.H. and Pavlath, G.K. (2010) Loss of nuclear poly(A)-binding protein 1 causes defects in myogenesis and mRNA biogenesis. Hum. Mol. Genet., 19, 1058–1065.

6. Iglesias, N., Tutucci, E., Gwizdek, C., Vinciguerra, P., Von Dach, E., Corbett, H., Dargemont, C. and Stutz, F. (2010) Ubiquitin-mediated mRNP dynamics and surveillance prior to budding yeast mRNA export. Genes Dev., 24, 1927–1938.

7. Kwiatek, L., Landry-Voyer, A.M., Latour, M., Yague-Sanz, C. and Bachand, F. (2023) PABPN1 prevents the nuclear export of an unspliced RNA with a constitutive transport element and controls human gene expression via intron retention. RNA, 29, 644–662.

8. Kühn, U. and Wahle, E. (2004) Structure and function of poly(A) binding proteins. Biochim. Biophys. Acta, 1678, 67–84.

9. Lima, S.A., Chipman, L.B., Nicholson, A.L., Chen, Y.H., Yee, B.A., Yeo, G.W., Coller, J. and Pasquinelli, A.E. (2017) Short poly(A) tails are a conserved feature of highly expressed genes. Nat. Struct. Mol. Biol., 24, 1057–1063.

10. Passmore, L.A. and Tang, T.T. (2021) The long and short of it. Elife, 10, e70757.

11. Wu, X., Liu, M., Downie, B., Liang, C., Ji, G., Li, Q.Q. and Hunt, A.G. (2011) Genome-wide landscape of polyadenylation in Arabidopsis provides evidence for extensive alternative polyadenylation. Proc. Natl. Acad. Sci. U.S.A., 108, 12533–12538.

12. Zhang, Y., Liu, L., Qiu, Q., Zhou, Q., Ding, J., Lu, Y. and Liu, P. (2021) Alternative polyadenylation: methods, mechanism, function, and role in cancer. J. Exp. Clin. Cancer Res. 40, 51.

13. Rodríguez-Molina, J.B. and Turtola, M. (2023) Birth of a poly(A) tail: mechanisms and control of mRNA polyadenylation. FEBS Open Bio., 13, 1140–1153.

14. Hunt, A.G. (2008) Messenger RNA 3’ end formation in plants. Curr. Top. Microbiol. Immunol., 326, 151–177.

15. Xing, D. and Li, Q.Q. (2011) Alternative polyadenylation and gene expression regulation in plants. Wiley Interdiscip. Rev. RNA, 2, 445–458.

16. Proudfoot, N. and Brownlee, G. (1976) 3’ non-coding region sequences in eukaryotic messenger RNA. Nature, 263, 211–214.

17. Loke, J.C., Stahlberg, E.A., Strenski, D.G., Haas, B.J., Wood, P.C. and Li, Q.Q. (2005) Compilation of mRNA polyadenylation signals in Arabidopsis revealed a new signal element and potential secondary structures. Plant Physiol., 138, 1457–1468.

18. Ye, C., Zhao, D., Ye, W., Wu, X., Ji, G., Li, Q.Q. and Lin, J. (2021) QuantifyPoly(A): reshaping alternative polyadenylation landscapes of eukaryotes with weighted density peak clustering. Brief. Bioinform., 22, bbab268.

19. Millevoi, S. and Vagner, S. (2010) Molecular mechanisms of eukaryotic pre-mRNA 3’ end processing regulation. Nucleic Acids. Res., 38, 2757–2774.

20. Bernardes, W.S. and Menossi, M. (2020) Plant 3’ Regulatory Regions From mRNA-Encoding Genes and Their Uses to Modulate Expression. Front. Plant Sci., 11, 1252.

21. Hunt, A.G., Xu, R., Addepalli, B., Rao, S., Forbes, K.P., Meeks, L.R., Xing, D., Mo, M., Zhao, H., Bandyopadhyay, A., Dampanaboina, L., Marion, A., Von Lanken, C. and Li, Q.Q. (2008) Arabidopsis mRNA polyadenylation machinery: comprehensive analysis of protein-protein interactions and gene expression profiling. BMC Genomics, 9, 220.

22. Hunt, A.G., Xing, D. and Li, Q.Q. (2012) Plant polyadenylation factors: conservation and variety in the polyadenylation complex in plants. BMC Genomics, 13, 641.

23. Zhang, X., Nomoto, M., Garcia-León, M., Takahashi, N., Kato, M., Yura, K., Umeda, M., Rubio, V., Tada, Y., Furumoto, T., Aoyama, T. and Tsuge, T. (2022) CFI 25 Subunit of Cleavage Factor I is Important for Maintaining the Diversity of 3’ UTR Lengths in Arabidopsis thaliana (L.) Heynh. Plant Cell Physiol., 63, 369–383.

24. Hardy, J.G. and Norbury, C.J. (2016) Cleavage factor Im (CFIm) as a regulator of alternative polyadenylation. Biochem. Soc. Trans., 44, 1051–1057.

25. Tian, B., Hu, J., Zhang, H. and Lutz, C.S. (2005) A large-scale analysis of mRNA polyadenylation of human and mouse genes. Nucleic Acids Res., 33, 201–212.

26. Hogg, J.R. and Goff, S.P. (2010) Upf1 senses 3’UTR length to potentiate mRNA decay. Cell, 143, 379–389.

27. Hoffman, Y., Bublik, D.R., Ugalde, A.P., Elkon, R., Biniashvili, T., Agami, R., Oren, M. and Pilpel, Y. (2016) 3’UTR Shortening Potentiates MicroRNA-Based Repression of Pro-differentiation Genes in Proliferating Human Cells. PLoS Genet., 12, e1005879. Erratum in: *PLoS Genet*., **12**, e1006254.

28. Chen, L.L. and Carmichael, G.G. (2009) Altered nuclear retention of mRNAs containing inverted repeats in human embryonic stem cells: functional role of a nuclear noncoding RNA. Mol. Cell, 35, 467–478.

29. An, J.J., Gharami, K., Liao, G.Y., Woo, N.H., Lau, A.G., Vanevski, F., Torre, E.R., Jones, K.R., Feng, Y., Lu, B. and Xu, B. (2008) Distinct role of long 3’ UTR BDNF mRNA in spine morphology and synaptic plasticity in hippocampal neurons. Cell, 134, 175–187.

30. Reid, D.W. and Nicchitta, C.V. (2015) Diversity and selectivity in mRNA translation on the endoplasmic reticulum. Nat. Rev. Mol. Cell Biol., 16, 221–231.

31. Alt, F.W., Bothwell, A.L., Knapp, M., Siden, E., Mather, E., Koshland, M. and Baltimore, D. (1980) Synthesis of secreted and membrane-bound immunoglobulin mu heavy chains is directed by mRNAs that differ at their 3’ ends. Cell, 20, 293–301.

32. Kim, S., Yamamoto, J., Chen, Y., Aida, M., Wada, T., Handa, H. and Yamaguchi, Y. (2010) Evidence that cleavage factor Im is a heterotetrameric protein complex controlling alternative polyadenylation. Genes Cells, 15, 1003–1013.

33. Gruber, A.R., Martin, G., Keller, W. and Zavolan, M. (2012) Cleavage factor Im is a key regulator of 3’ UTR length. RNA Biol., 9, 1405–1412.

34. Yang, Q., Gilmartin, G.M. and Doublié, S. (2010) Structural basis of UGUA recognition by the Nudix protein CFI(m)25 and implications for a regulatory role in mRNA 3’ processing. Proc. Natl. Acad. Sci. U.S.A., 107, 10062–10067.

35. Yang, Q., Coseno, M., Gilmartin, G.M. and Doublié, S. (2011) Crystal structure of a human cleavage factor CFI(m)25/CFI(m)68/RNA complex provides an insight into poly(A) site recognition and RNA looping. Structure, 19, 368–377.

36. Martin, G., Gruber, A.R., Keller, W. and Zavolan, M. (2012) Genome-wide analysis of pre-mRNA 3’ end processing reveals a decisive role of human cleavage factor I in the regulation of 3’ UTR length. Cell Rep., 1, 753–763.

37. Zhu, Y., Wang, X., Forouzmand, E., Jeong, J., Qiao, F., Sowd, G.A., Engelman, A.N., Xie, X., Hertel, K.J. and Shi, Y. (2018) Molecular Mechanisms for CFIm-Mediated Regulation of mRNA Alternative Polyadenylation. Mol Cell, 69, 62–74.

38. Li, N., Cai, Y., Zou, M., Zhou, J., Zhang, L., Zhou, L., Xiang, W., Cui, Y. and Li, H. (2023) CFIm-mediated alternative polyadenylation safeguards the development of mammalian pre-implantation embryos. Stem Cell Reports, 18, 81–96.

39. Tseng, H.W., Mota-Sydor, A., Leventis, R., Jovanovic, P., Topisirovic, I. and Duchaine, T.F. (2022) Distinct, opposing functions for CFIm59 and CFIm68 in mRNA alternative polyadenylation of Pten and in the PI3K/Akt signalling cascade. Nucleic Acids Res., 50, 9397–9412.

40. Alonso, J.M., Stepanova, A.N., Leisse, T.J., Kim, C.J., Chen, H., Shinn, P., Stevenson, D.K., Zimmerman, J., Barajas, P., Cheuk, R., Gadrinab, C., Heller, C., Jeske, A., Koesema, E., Meyers, C.C., Parker, H., Prednis, L., Ansari, Y., Choy, N., Deen, H., Geralt, M., Hazari, N., Hom, E., Karnes, M., Mulholland, C., Ndubaku, R., Schmidt, I., Guzman, P., Aguilar-Henonin, L., Schmid, M., Weigel, D., Carter, D.E., Marchand, T., Risseeuw, E., Brogden, D., Zeko, A., Crosby, W.L., Berry, C.C., Ecker, J.R. (2003) Genome-wide insertional mutagenesis of Arabidopsis thaliana. Science, 301, 653–657.

41. Kleinboelting, N., Huep, G., Kloetgen, A., Viehoever, P. and Weisshaar, B. (2012) GABI-Kat SimpleSearch: new features of the Arabidopsis thaliana T-DNA mutant database. Nucleic Acids Res., 40, D1211–D1215.

42. Clough, S.J. and Bent, A.F. (1998) Floral dip: a simplified method for Agrobacterium-mediated transformation of Arabidopsis thaliana. Plant J., 16, 735–743.

43. Grefen, C., Donald, N., Hashimoto, K., Kudla, J., Schumacher, K. and Blatt, M.R. (2010) A ubiquitin-10 promoter-based vector set for fluorescent protein tagging facilitates temporal stability and native protein distribution in transient and stable expression studies. Plant J., 64, 355–365.

44. Tzfira, T., Tian, G.W., Lacroix, B., Vyas, S., Li, J., Leitner-Dagan, Y., Krichevsky, A., Taylor, T., Vainstein, A. and Citovsky, V. (2005) pSAT vectors: a modular series of plasmids for autofluorescent protein tagging and expression of multiple genes in plants. Plant Mol. Biol., 57, 503–516.

45. Merzlyak, E.M., Goedhart, J., Shcherbo, D., Bulina, M.E., Shcheglov, A.S., Fradkov, A.F., Gaintzeva, A., Lukyanov, K.A., Lukyanov, S., Gadella, T.W. and Chudakov, D.M. (2007) Bright monomeric red fluorescent protein with an extended fluorescence lifetime. Nat. Methods, 4, 555–557.

46. Imajuku, Y., Ohashi, Y., Aoyama, T., Goto, K. and Oka, A. (2001) An upstream region of the Arabidopsis thaliana CDKA;1 (CDC2aAt) gene directs transcription during trichome development. Plant Mol. Biol., 46, 205–213.

47. Alexander, M.P. (1969) Differential staining of aborted and nonaborted pollen. Stain Technol. 44, 117–122.

48. Kuhn, L., Vincent, T., Hammann, P. and Zuber, H. (2023) Exploring Protein Interactome Data with IPinquiry: Statistical Analysis and Data Visualization by Spectral Counts. Methods Mol. Biol., 2426, 243–265.

49. Robinson, M.D., McCarthy, D.J. and Smyth, G.K. (2010) edgeR: a Bioconductor package for differential expression analysis of digital gene expression data. Bioinformatics, 26, 139–140.

50. Love, M.I., Huber, W. and Anders, S. (2014) Moderated estimation of fold change and dispersion for RNA-seq data with DESeq2. Genome Biol., 15, 550.

51. Bajczyk, M., Lange, H., Bielewicz, D., Szewc, L., Bhat, S.S., Dolata, J., Kuhn, L., Szweykowska-Kulinska, Z., Gagliardi, D. and Jarmolowski, A. (2020) SERRATE interacts with the nuclear exosome targeting (NEXT) complex to degrade primary miRNA precursors in Arabidopsis. Nucleic Acids Res., 48, 6839–6854.

52. Pati, P.K., Ma, L. and Hunt, A.G. (2015) Genome-wide determination of poly(A) site choice in plants. Methods Mol. Biol., 1255, 159–174.

53. Smith, T., Heger, A. and Sudbery, I. (2017) UMI-tools: modeling sequencing errors in Unique Molecular Identifiers to improve quantification accuracy. Genome Res., 27, 491–499.

54. Martin, M. (2011) Cutadapt removes adapter sequences from high-throughput sequencing reads. EMBnet J., 17, 10–12.

55. Kim, D., Paggi, J.M., Park, C., Bennett, C. and Salzberg, S.L. (2019) Graph-based genome alignment and genotyping with HISAT2 and HISAT-genotype. Nat. Biotechnol., 37, 907–915.

56. Ye, C., Zhao, D., Ye, W., Wu, X., Ji, G., Li, Q.Q. and Lin, J. (2021) QuantifyPoly(A): reshaping alternative polyadenylation landscapes of eukaryotes with weighted density peak clustering. Brief Bioinform. 22, bbab268.

57. Quinlan, A.R. and Hall, I.M. (2010) BEDTools: a flexible suite of utilities for comparing genomic features. Bioinformatics, 26, 841–842.

58. Li, H., Handsaker, B., Wysoker, A., Fennell, T., Ruan, J., Homer, N., Marth, G., Abecasis, G., Durbin, R. and 1000 Genome Project Data Processing Subgroup, (2009) The Sequence Alignment/Map format and SAMtools. Bioinformatics, 25, 2078–2079.

59. Robinson, J.T., Thorvaldsdóttir, H., Winckler, W., Guttman, M., Lander, E.S., Getz, G. and Mesirov, J.P. (2011). Integrative genomics viewer. Nat. Biotechnol., 29, 24–26.

60. Smyth, D.R., Bowman, J.L. and Meyerowitz. (1990) Early flower development in Arabidopsis. Plant Cell, 2, 755–767.

61. Kosugi, S., Hasebe, M., Tomita, M. and Yanagawa, H. (2009) Systematic identification of cell cycle-dependent yeast nucleocytoplasmic shuttling proteins by prediction of composite motifs. Proc. Natl. Acad. Sci. U.S.A., 25, 10171–10176.

62. Venkataraman, K., Brown, K.M. and Gilmartin, G.M. (2005) Analysis of a noncanonical poly(A) site reveals a tripartite mechanism for vertebrate poly(A) site recognition. Genes Dev., 19, 1315–1327.

63. Addepalli, B. and Hunt, A.G. (2007) A novel endonuclease activity associated with the Arabidopsis ortholog of the 30-kDa subunit of cleavage and polyadenylation specificity factor. Nucleic Acids Res., 35, 4453–4463.

64. So, B.R., Di, C., Cai, Z., Venters, C.C., Guo, J., Oh, J.M., Arai, C. and Dreyfuss, G. (2019) A Complex of U1 snRNP with Cleavage and Polyadenylation Factors Controls Telescripting, Regulating mRNA Transcription in Human Cells. Mol. Cell, 76, 590–599.

65. Zhou, L., Li, K. and Hunt, A.G. (2024) Natural variation in the plant polyadenylation complex. Front. Plant Sci., 14, 1303398.

66. SoLuchsinger, C., Lee, K., Mardones, G.A., KewalRamani, V.N. and Diaz-Griffero, F. (2024) Formation of nuclear CPSF6/CPSF5 biomolecular condensates upon HIV-1 entry into the nucleus is important for productive infection. Sci. Rep., 13, 10974.

67. Ay, S. and Di Nunzio, F. (2023) HIV-Induced CPSF6 Condensates. J. Mol. Biol., 435, 168094.

68. Lee, J.H., Yi, L., Li, J, Schweitzer, K., Borgmann, M, Naumann, M. and Wu, H. (2013) Crystal structure and versatile functional roles of the COP9 signalosome subunit 1. Proc. Natl. Acad. Sci. U.S.A. 110, 11845–11850.

69. Ramírez-Aportela, E., López-Blanco, J.R. and Chacón, P. (2016) FRODOCK 2.0: fast protein-protein docking server. Bioinformatics, 32, 2386–2388.

