## Supplementary Figures for "Plant Cleavage Factor I complex is essential for precise cleavage and polyadenylation site determination"

```
HsCFI 68     1 -----------------------------MADG--VDHIDIYADV--GEEFNQEAEYG--
HsCFI 59     1 -----------------------------MSEG--VDLIDIYAD----EEFNQDPEFN--
AtCFI 68     1 -MDEGDGRDQMDQFHQN-EAISAVADDGFMAEEEDDDYEDLYNDVNVGEGFLQSMKKNDE
AtCFI 59     1 MTEENDYGGNQKILHQGSGTIPALADEELMGD--DDEYDDLYSDVNVGESFFQAHNQPQP

HsCFI 68    26 ------------------------------------------------GHDQIDLYDDVI
HsCFI 59    24 ------------------------------------------------NTDQIDLYDDVL
AtCFI 68    59 AGSRNEEKEKVNMEEEDRVEPVLGEAEVSISIPGLVGESVEKEAEAESGGGGSGSGTDVV
AtCFI 59    59 PAQVNTSNASLQAQN----SHVAAEPRMGIVSGGTVEGKYRNDGGHN-GISGPDTRSDVY

HsCFI 68    38 SPSANN------------------------------GDAPEDRDYMDTL-----------
HsCFI 59    36 TATSQ---------------------------------PSDDRSSSTEP-----------
AtCFI 68   119 VASSGYGAQEVKVSDVSQEIPGGIGTGTGGGLRVELGQASNRANDLEAPRGNNISQGLLP
AtCFI 59   114 PQASSFGAKGLNIDIQSNKIAQ---QGSTTVVLNNHGFSGNAVNVPEMPVHNSYG-----

HsCFI 68    57 --PPTVGDDVG---KGAAPNVVYTYTG--------------------------KRIALYI
HsCFI 59    52 --PPPVRQEPSPKPNNKTPAILYTYSGLR-----------------------NRRAAVYV
AtCFI 68   179 PPPVLGNNENLMRPVMGNVNGGIPPGPGSNMVGNGANIAMPGVVGGGTGGGGGGAFLFV
AtCFI 59   166 -APPQGAQQIPVSQMSVNPNVMMNKSPTQSFVVD------------------NGNTMLFV

HsCFI 68    86 GNLTWWTTDEDLTEAVHSLGVNDILEIKFFENRANGQSKGFALVGVGSEASSKKLMDLLP
HsCFI 59    87 GSFSWWTTDQQLIQVIRSIGVYDVVELKFAENRANGQSKGYAEVVVASENSVHKLLELLP
AtCFI 68   239 GDLHWWTTDAELEAELCKYGA--VKEVKFFDEKASGKSKGYCQVEFYDPVAASACKDALN
AtCFI 59   207 GELHWWTTDAEIESVLSQYGR--VKEIKFFDERVSGKSKGYCQVEFYDSAAAAACKEGMN

HsCFI 68   146 KRELHGQNPVVTPCNKQFLSQFEMQSRKTTQSGQMSGEGK------AGPPGGSSRAAFPQ
HsCFI 59   147 GKVLNGEKVDVRPATRQNLSQFEAQARKRECVRVPRG---------GIPPRAHSRDSS-D
AtCFI 68   297 GYPFNGRPCVVEYASPYSVKRMGEAQVNRTQQAQSVIAQAKRGGPADPPSKPLVANNNNN
AtCFI 59   265 GFIFNGKACVVAFASPETLKQMGANFTGRNQGQNQIQNRR-------PLNEGMGRGNNNN

HsCFI 68   200 GGRGRGRFPGAVPGGDRFPGPAGPGGPPPPFPAGQTPPRPPLGPPGPPGPPGPPPPGQVL
HsCFI 59   197 SADGR-----ATPSENLVPSSARVDKPPSVLPYFNRPP-SALPLMGLPPPPIPPPP----
AtCFI 68   357 NNNNAIGGNFQGGENRGFGRGNWGRGNAQGMGGRG--PGGPMRNR-PNGMGGRGLMGNGG
AtCFI 59   318 NNMNTQ----NGDGGRNYGRGGFARG-GQGMGNRGGAWGGAMRGRGVNNMASG—SGAGP

HsCFI 68   260 PPPLAGPPNRGDRPPPPVLFPGQPFGQPPLGPLPPGPPPPVPGYGPPPGPPPPQQGPPPP
HsCFI 59   247 --PLS-----------------SSFGVPP----------------PPPGIHYQHLMPPPP
AtCFI 68   414 FGQGMGTGPPMNMMHQP-MMGQG-FEQAFGGPM---------ARMGGYGGFPGAPGPQFP
AtCFI 59   371 YGPGLAGPAFGGMMHPQGMMGAGGFDPTFMG------------RGAGYGGYS---GIAYP

HsCFI 68   320 PGPFPPRPPGPLGPPLTLAPPPHLPGPPPGAPPPAPHVNPAFFPPPTNSGMPTSDSRGPP
HsCFI 59   272 ------------------RLPPHLAVPPPGAIPPALHLNPAFFPPPN-------ATVGPP
AtCFI 68   463 G-----------------LLSSFPPVGGVGLPGVAPHVNPAFFGRGM-----PMNGMGMM
AtCFI 59   416 G-----------------MPHSYPGVNAMGMVGIAPHVNPAFFG----------TGMGTM

HsCFI 68   380 PTDPYGRPPPYDRGDYGPPGREMDTA------------RTPLSEAEFEEIMNRNRAISSS
HsCFI 59   307 PDTYMKASAPYN--HHGS--RDSGPP------------PSTVSEAEFEDIMKRNRAISSS
AtCFI 68   501 PNAGVDGGHNMGMWDPNSGGWGAGEDLGSGRAAESSYGEEAASDHQYGEVNHERGARPN-
AtCFI 59   449 GSSGMNGVHAAAMWSEANGGGGE---------------EGGSEYGGYEDETQEKEEKPS-
```

Proline-rich

**Supplementary Figure S1.** AtCFI 59 and AtCFI 68 harbored the conserved RRM domain, proline-rich domain, and RS/RD/RE region. AtCFI 59 and AtCFI 68 consisted of 573 aa and 710 aa, with a predicted molecular mass of 61.4 KDa and 76.4 KDa, respectively. At the amino acid level, identity between AtCFI 59 and AtCFI 68 was 34.99%. The GAR motif is indicated by a red box. The RRM domain is indicated by a red line. The proline rich region is indicated by a yellow line. The RS/RD/RE region is indicated by a black line. The amino acid sequences encoded by homologous proteins in Arabidopsis and human were aligned using the CLUSTALW (2.1) and BoxShade (3.2). The black boxes mark residues identical in more than two proteins, while the grey boxes mark residues of similarity.

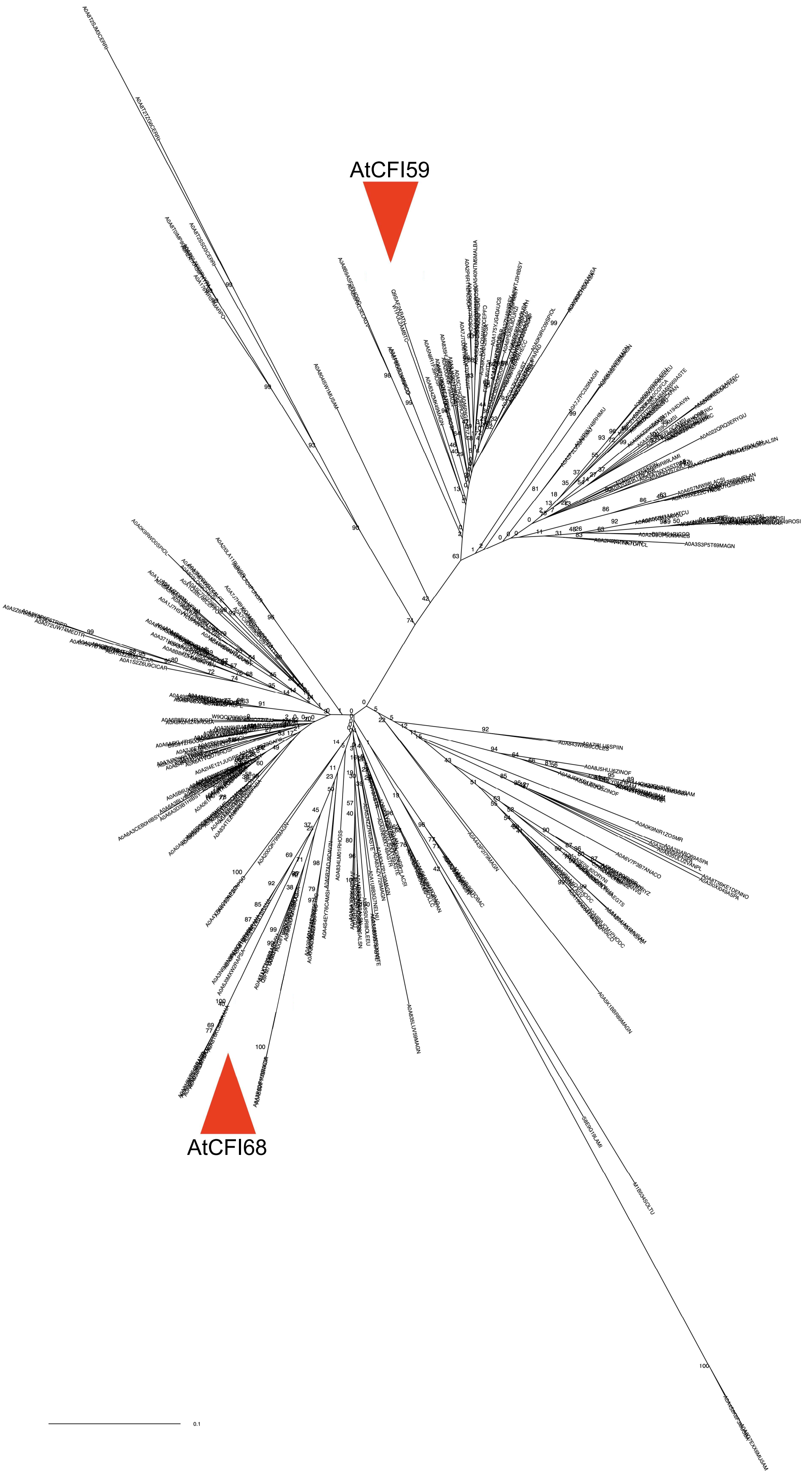


**Supplementary Figure S2.** AtCFI59 and AtCFI68 is assigned to two different clades in the phylogenic tree of CFI59, CFI68 and their homologs in higher plants. 6,000 sequences were selected from 26,000 related sequences to extract approximately 100 amino acids of RRM. Mr. Baves software was used for calculation.

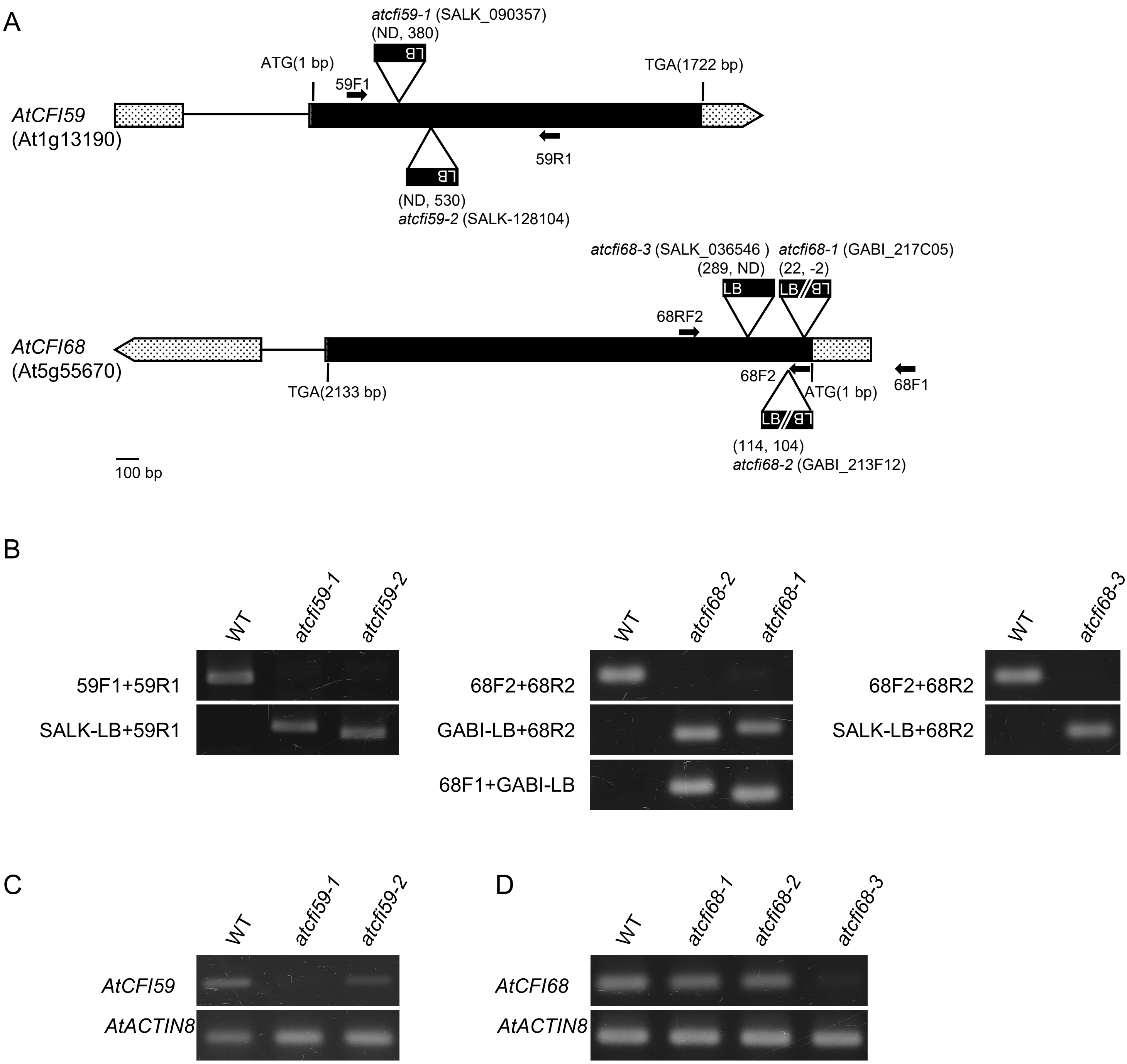


**Supplementary Figure S3.** Establishing loss of function mutants of *AtCFI59* and *AtCFI68*. **(A)** Schematic diagram of *AtCFI59* and *AtCFI68* genomic structure with T-DNA insertion sites. Exons are shown as black boxes, introns as black lines, 5' UTR as gray boxes, 3' UTR as gray arrowed boxes, and intergenic region as broken lines. Black arrows indicate positions of primers used for PCR-based genotyping in (B). T-DNA insertion sites were determined by sequencing flanking regions. Numbers in parenthesis correspond to the flanking genomic position of the T-DNA insertion. ND: not determined, LB: left border. **(B)** PCR-based genotyping for mutant alleles of *atcfi59* and *atcfi68*. Loss of function plants, carrying homozygote single T-DNA copy were, established and analyzed using primers shown in Supplementary Table S1. SALK-LB primer was used for*, atcfi59-1*, *atcfi59-2*, and *atcfi68-3*, which is specific for the pROK2 T-DNA vector. GABI-LB was used for *atcfi68-1* and *atcfi68-2*, which is specific for the pAC161 T-DNA vector (https://www.osu.edu). **(C)** Semi-quantitative RT-PCR analysis of *AtCFI59* transcripts in two independent lines of *atcfi59*. Transcripts of *AtCFI59* were not detected in *atcfi59-1*, indicating that *atcfi59-1* was a null allele for *AtCFI 59*. **(D)** Semi-quantitative RT-PCR analysis of *AtCFI68* transcripts in three independent *atcfi68* lines. Note that transcripts of *AtCFI68* were not detectable in *atcfi68-3*, indicating that *atcfi68-3* was a null allele for *AtCFI68.*

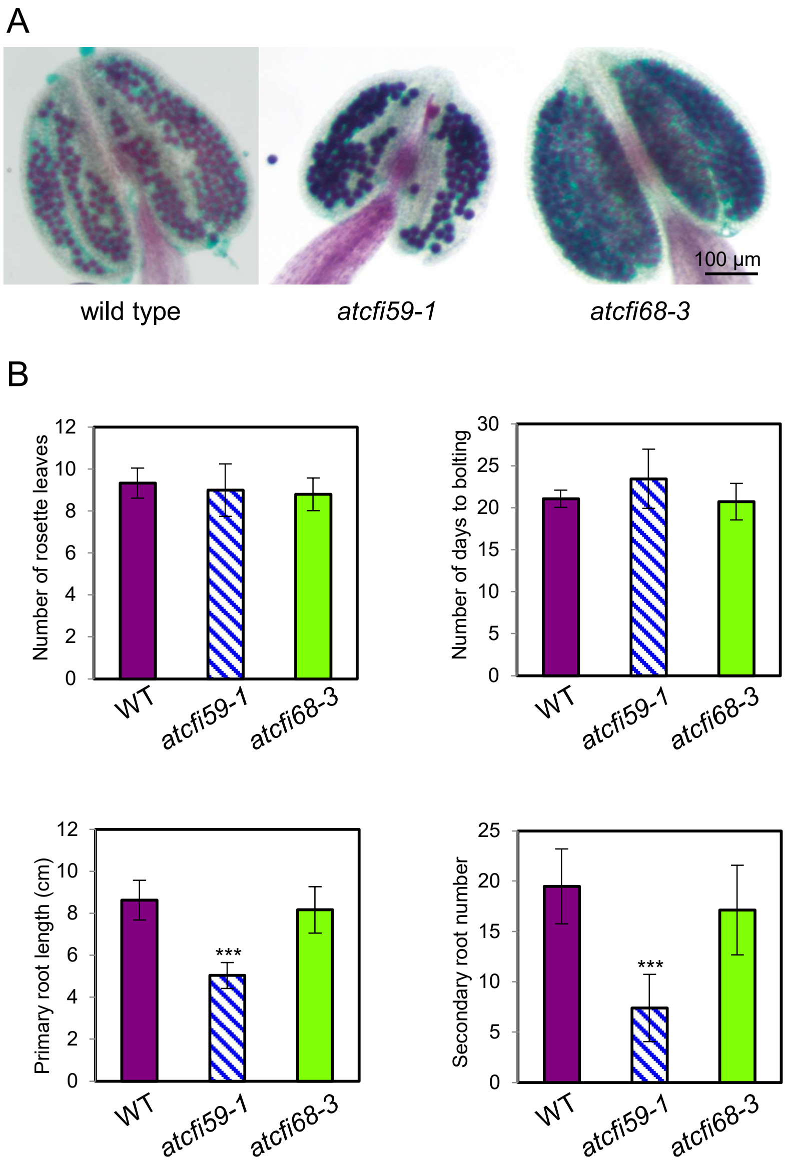


**Supplementary Figure S4. (A)** Alexander's staining showed no obvious difference in the viability of the pollen between *atcfi59-1*, *atcfi68-3*, and wild type. Note that *atcfi59-1* had smaller anther with less pollen granules. **(B)** The flowering time of *atcfi59-1* and *atcfi68-3* were indistinguishable from that of wild type. *atcfi59-1* had significantly shorter primary roots and fewer numbers of lateral roots, while *atcfi68-3* showed no obvious difference with the wild type. Subjected samples: 15 DAS for root analyses (B). Student's t-test: ***P < 0.001; **P < 0.01; *P < 0.05. Lengths of scale bars are noted in the figure.

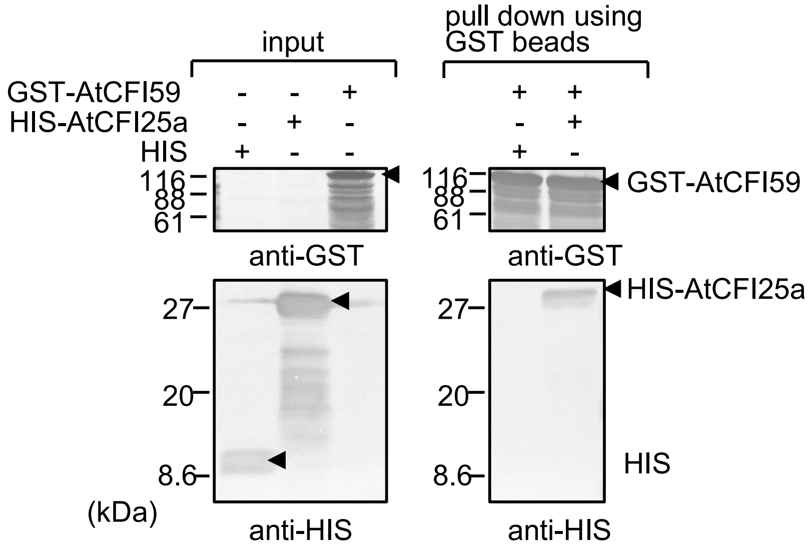


**Supplementary Figure S5.** AtCFI25a interacts with putative AtCFI subunits. Western blot analyses verified recombinant AtCFI25a bound AtCFI59. AtCFI25a was fused to HIS tag, while AtCFI59 was fused to GST-tag. Proteins were detected using anti-GST antibody following SDS-PAGE and membrane transfer. Pull-down conditions are noted at the top of the panel. Arrowheads indicate proteins of interest. Proteins were detected using anti-GST antibody following SDS-PAGE and membrane transfer.

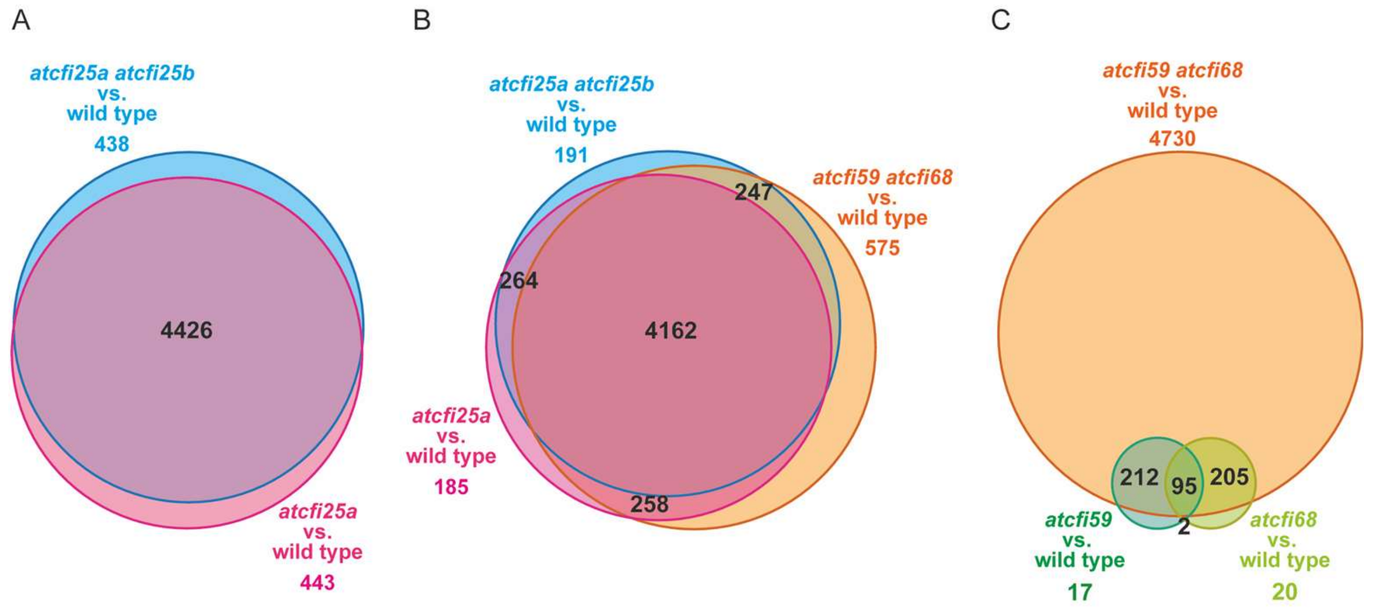


**Supplemental Figure S6.** Relationship of global polyadenylation changes among *atcfI* mutants in comparison to wild type. Venn diagrams of differential polyadenylation usage in groups of: **(A)** *atcfi25a* and *atcfi25a atcfi25b* double mutant, **(B)** *atcfi25a*, *atcfi25a atcfi25b* double mutant, and *atcfi59 atcfi68* double mutant, **(C)** *atcfi59* and *atcfi68*, **(D)** *atcfi59*, *atcfi68*, and *atcfi59 atcfi68* double mutant.

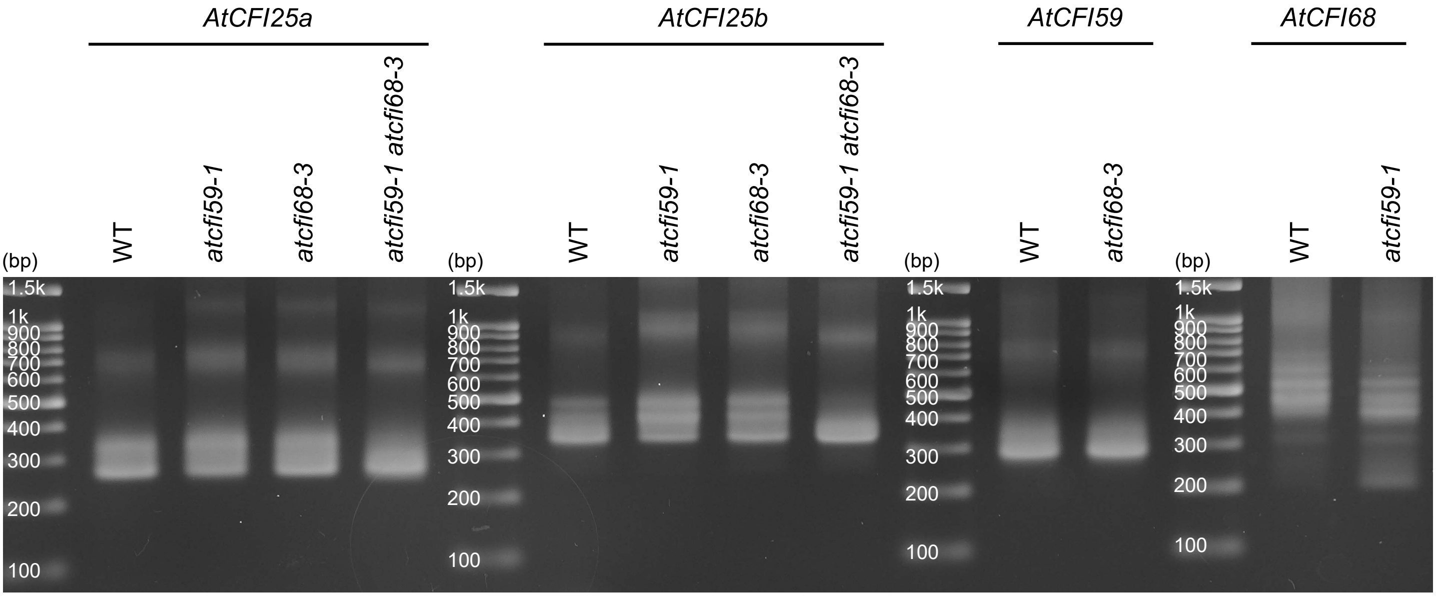


**Supplementary Figure S7.** Comparison of cleavage and polyadenylation sites of wild type, *atcfi59-1*, *atcfi68-3*, and *atcfi59-1 atcfi68-3* double mutant for genes encoding putative CFI subunits. 3' RACE amplifications of *AtCFI25a*, *AtCFI25b*, *AtCFI59*, and *AtCFI68* are shown as comparable gel patterns on agarose gel after electrophoresis. Note that the 3' UTR length pattern is less diverse in *atcfi59-1 atcfi68-3* double mutant, compared to wild type. Subjected samples: 7 DAS seedlings.

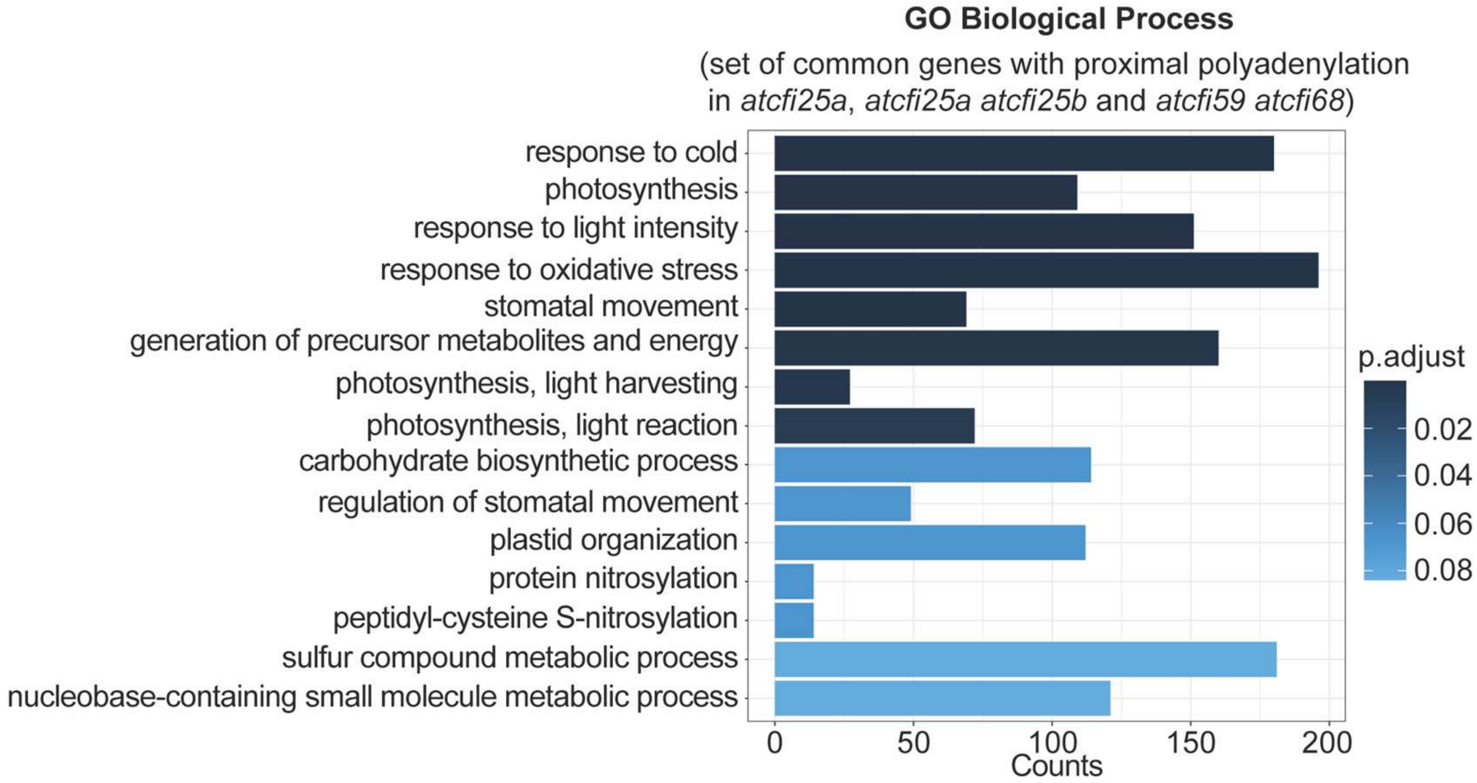


**Supplemental Figure S8.** Gene Ontology (GO) enrichment analysis of the common part between differentially polyadenylated genes in *atcfi25a*, *atcfi25a atcfi25b* double mutant, and *atcfi59 atcfi68* double mutant (common part extracted from the Venn diagram from the Fig. 8B). Bar chart showing the top 15 GO terms for biological processes. The fold enrichment is defined as the ratio of genes identified with the QuantifyPoly(A) tool that are annotated in a particular biological process to the number of all genes that are annotated in this biological process [Wu *et al*., 2021]

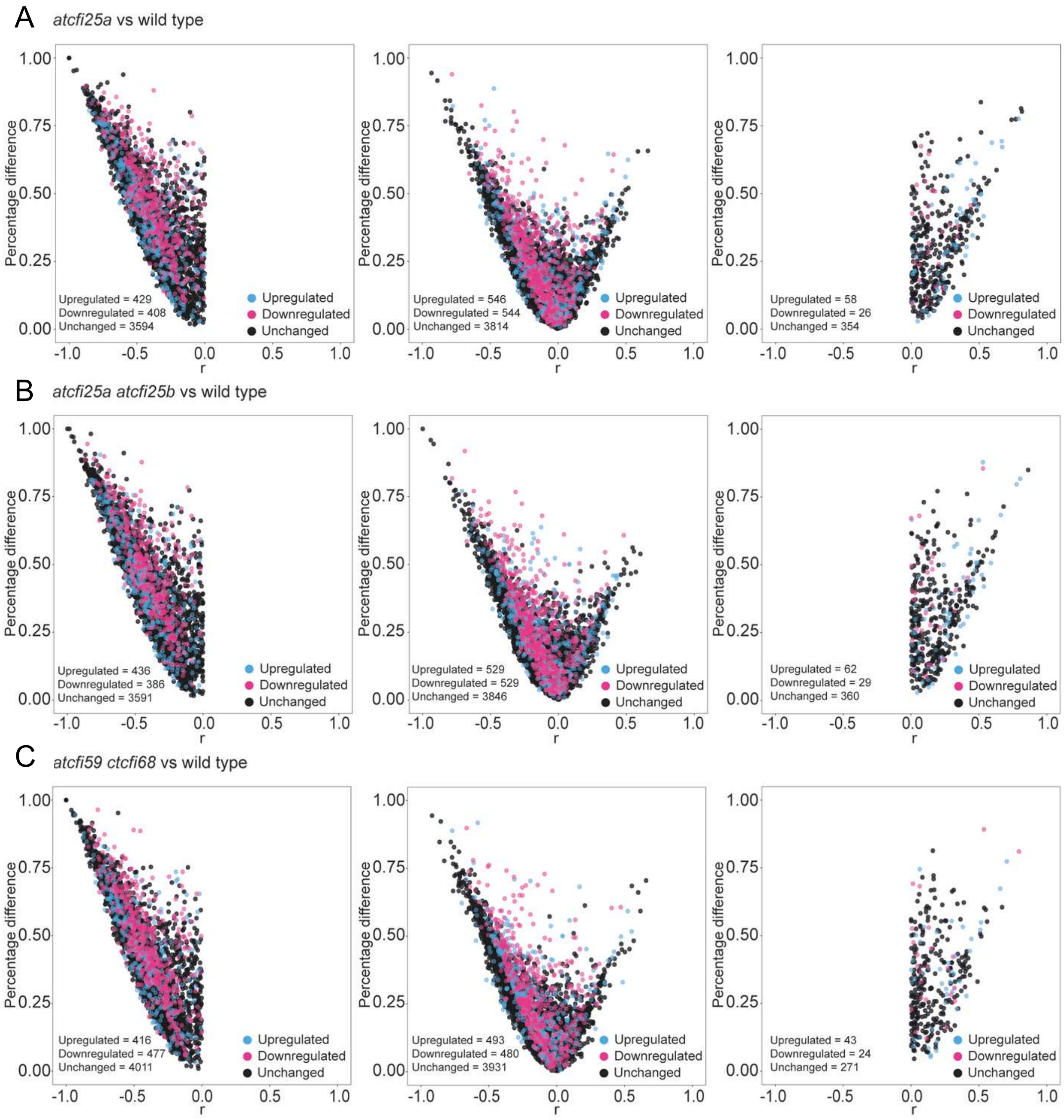


**Supplementary Figure S9.** Analysis of the association of changes in polyadenylation site selection with changes in expression levels. Information on expression levels was superimposed on the earlier information on the number of changes in polyadenylation site selection in individual mutants of the Arabidopsis CFI complex. Individual volcano plots (Fig. 8) of *atcfi25a* vs wt **(A)**, *atcfi25a atcfi25b* vs wt **(B)**, and *atcfi59 atcfi68* vs wt **(C)** were separated for changes in proximal polyadenylation site selection, distal polyadenylation site selection and for not statistically significant. Subsequently, information on the level of expression was added: transcripts with increased expression levels were marked in blue, transcripts with decreased expression levels were marked in magenta and those transcripts whose level did not change were marked in black. The analysis did not reveal any global trend regarding the effect of polyadenylation site selection on expression levels. A similar level of down-regulation and up-regulation of transcripts is seen in all cases analysed. For DEseq2 analysis fold change was set for 2 and false discovery rate was set for 0.05.

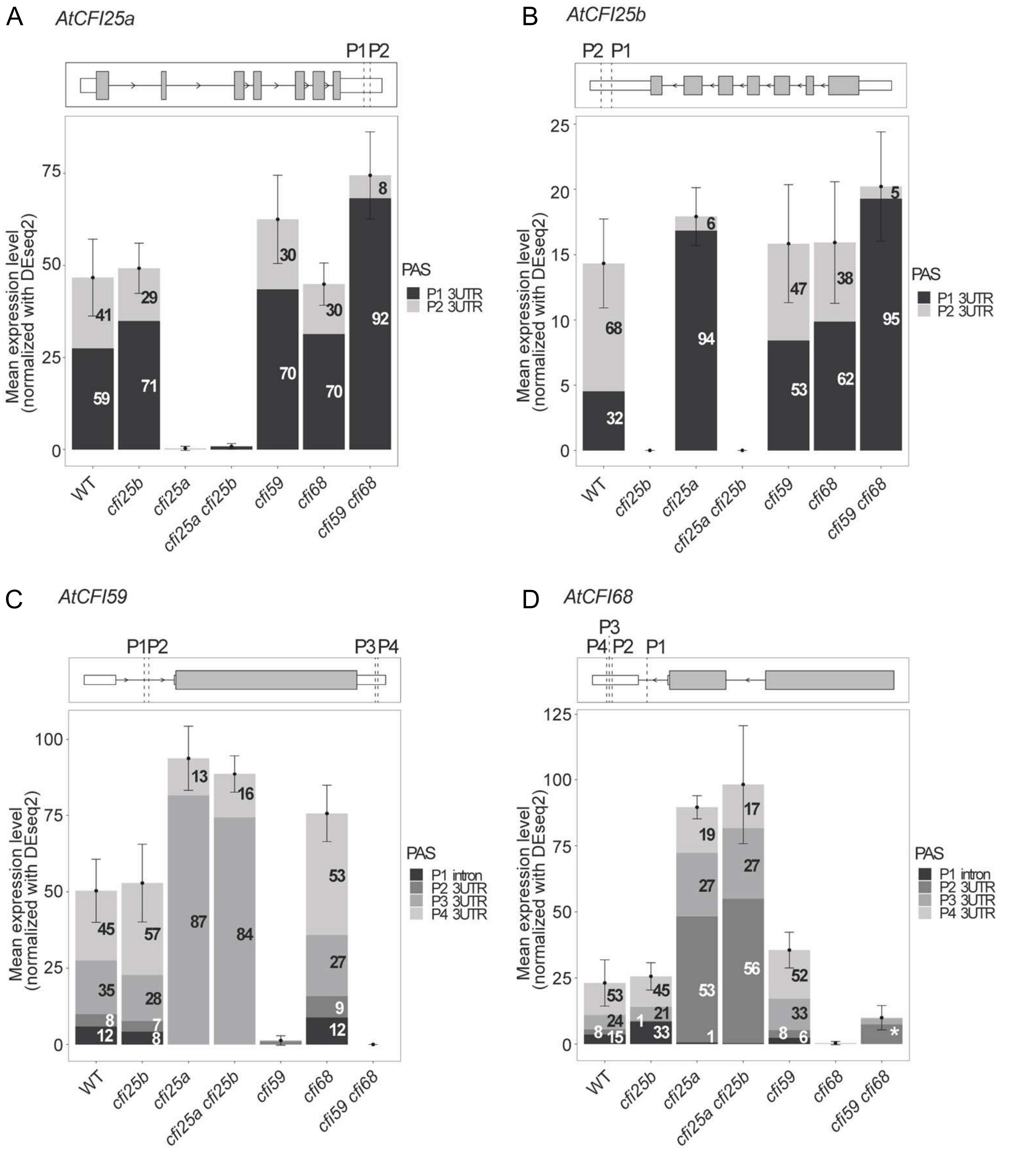


**Supplementary Figure S10.** Analysis of the expression level of *AtCFI25a* **(A)**, *AtCFI25b* **(B)**, *AtCFI59* **(C)**, and *AtCFI68* **(D)** combined with the polyadenylation site usage in the genetic background of different *cfi* mutants. The upper part of figures shows a schematic representation of each analysed gene with all identified polyadenylation sites marked. White rectangles represent UTRs, grey rectangles represent exons, black lines represent introns, arrowheads indicate the direction of transcription. Capital P with number marks the position of all identified polyadenylation sites (numbering starts from the transcription start site). The Y-axis shows the mean normalised counts, represented by the sum of all PAT-seq reads of the gene and normalised by DEseq2, the X-axis shows the investigated genotypes. The frequency of selection of a particular polyadenylation site is expressed as a percentage and indicated by numbers and the colour of each bar. The sum of all identified polyadenylation sites is 100%. A white asterisk marks the transcript of unknown origin in D. For an unknown reason, an increased number of reads assigned to the *AtCFI68* gene region has been observed in the *atcfi59 atcfi68* double mutant, although we do not see such reads in the *atcfi68* mutant. This may be related to the nature of T-DNA insertion mutants. T-DNA insertion may interrupt the production of the correct transcript, but it may not prevent transcription as such. In this case, we can observe the level of such aberrant transcripts.

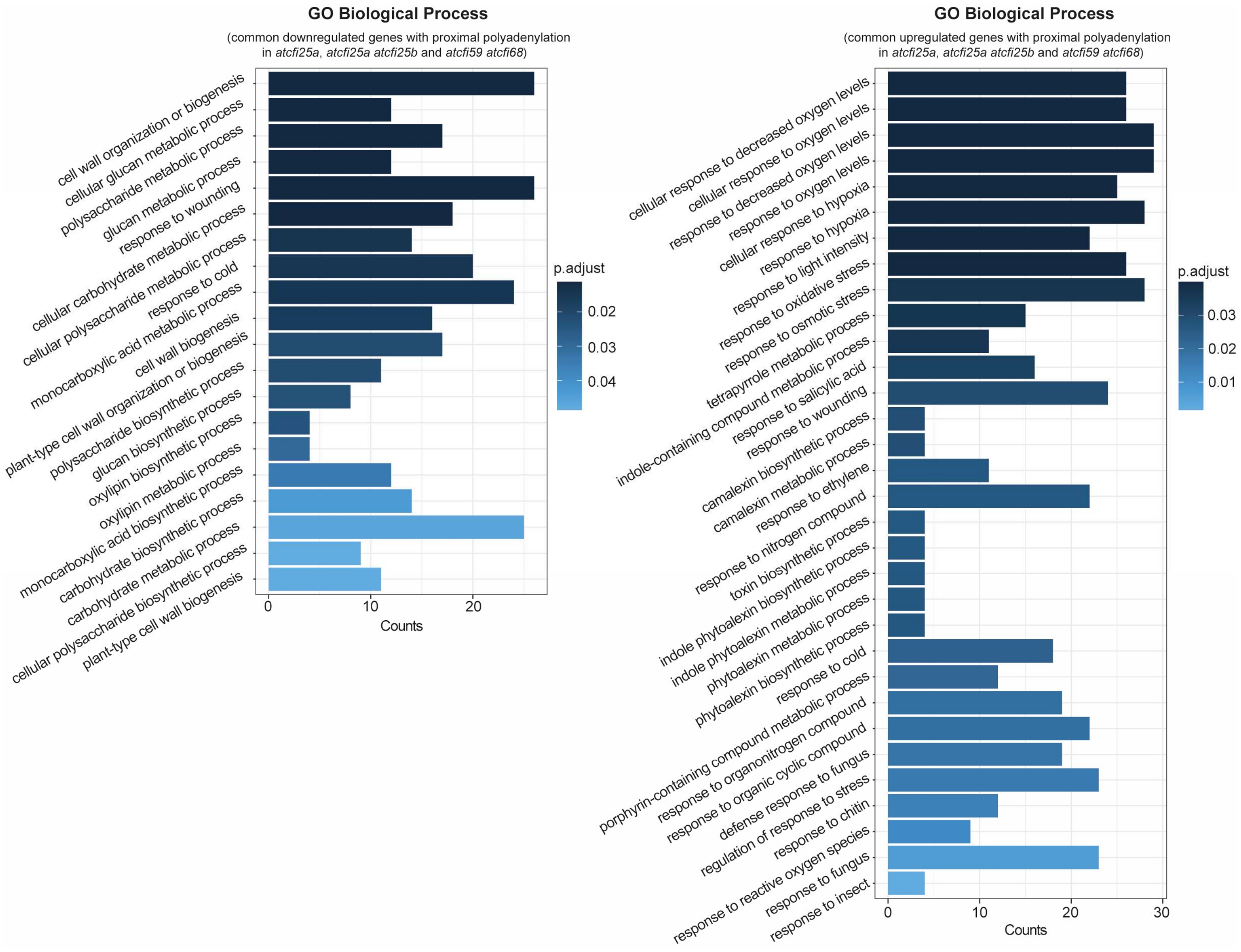


**Supplementary Figure S11.** Gene Ontology analysis of the group of genes identified as down- or upregulated in the common group of genes with proximal polyadenylation between *atcfi25a*, the *atcfi25a atcfi25b* double mutant, and the *atcfi59 atcfi68* double mutant (Supplementary Figure 10). Bar chart showing statistically significant hits for biological processes. The fold enrichment is defined as the ratio of down- or upregulated genes identified using the DEseq2 tool that are annotated in a particular biological process to the number of all genes annotated in that biological process [Wu *et al*. 2021].

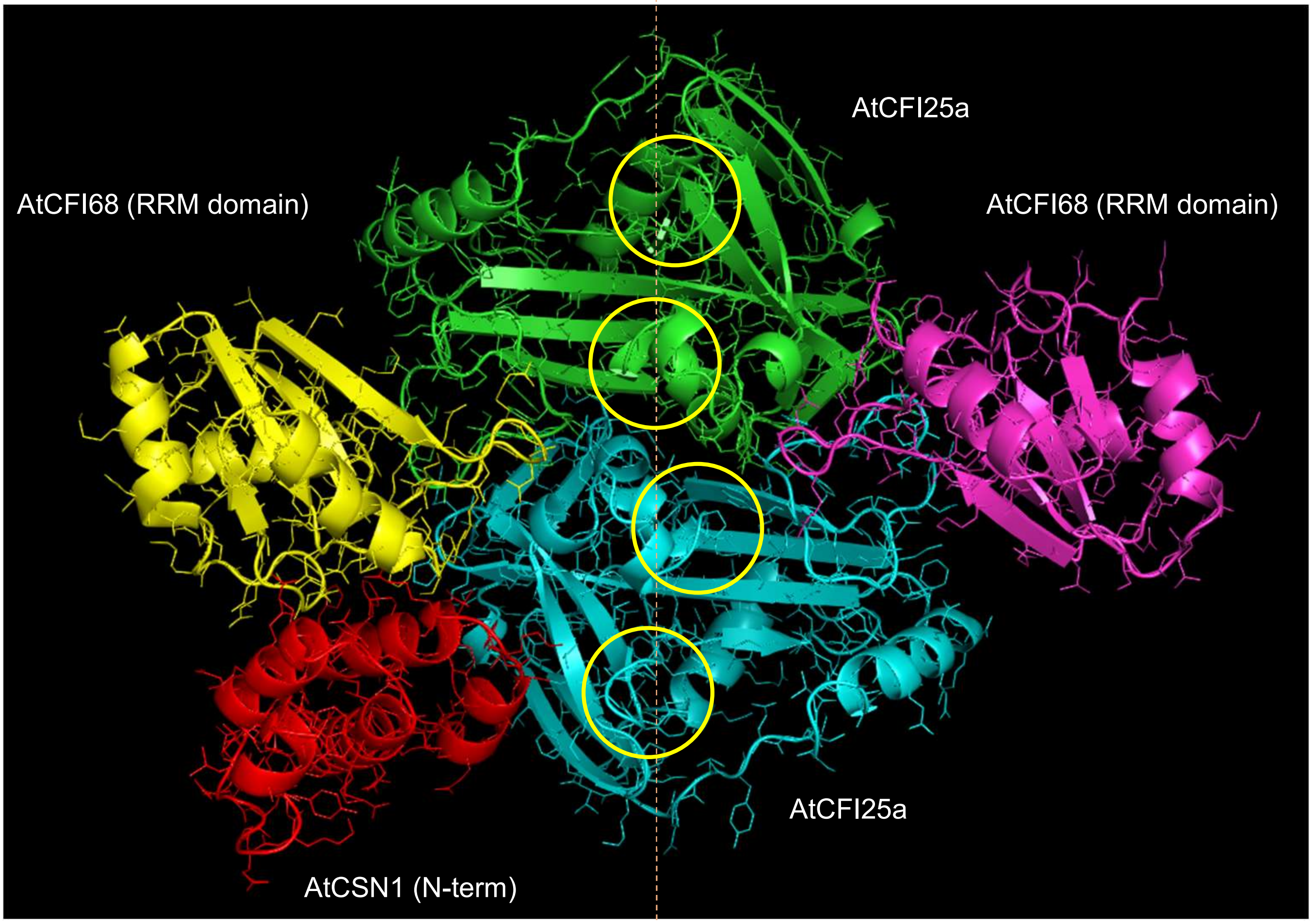


**Supplementary Figure S12.**
Comparative modelling of two AtCFI68 (RRM domains) with two AtCFI25a. The three-dimensional structure of AtCSN1 has been solved by [Lee *et al*. 2013]. The N-terminal region (1-110) of AtCNS1 was extracted from the coordinate stored in D chain of PDB ID 4lct. The three-dimensional structure of mammalian CFI68 binding RNA has been solved by analyzing the crystal structure [Yang *et al*. 2010, Yang *et al*. 2011]. Putative RNA binding sites of the AtCFI25a is illustrated as yellow circles superimposing the mammalian analyses. (Models were built and the model with the best DOPE energy score was selected. AtCFI25a, AtCFI68, and AtCSN1 interaction domain were docked on FRODOCK [Ramirez-Aportela,E., et al. 2016].
