## Supplementary Table S1 and S2 for "Plant Cleavage Factor I complex is essential for precise cleavage and polyadenylation site determination"

**Supplementary Table S1.**
Primers used in this study.

| Template | Sequence (5' - 3') | Main purpose | Primer name |
|---|---|---|---|
| AtCFI59 | AGCTGCTTCATAAATAATTCACCAG | Construction in binary vectors for GUS experiments | 59-promoter-F |
| AtCFI59 | TGAATTTATACAAACTGGCCAGAAA | Construction in binary vectors for GUS experiments | 59-promoter-R |
| AtCFI68 | GATTTTGCCGGCGGATGTAAAGT | Construction in binary vectors for GUS experiments | 68-promoter-F |
| AtCFI68 | CTTAAAAGACTCTTCCTTGTGAT | Construction in binary vectors for GUS experiments | 68-promoter-R |
| AtCFI68 | GGAGGAGGAAGATAGGGTTG | RT-PCR transcript detection | 68F3 |
| AtCFI68 | TATTCCCTCTAGGTGCCTCG | RT-PCR transcript detection | 68R1 |
| AtCFI59 | GATGAACAAAAGCCCTACGC | RT-PCR transcript detection | 59F2 |
| AtCFI59 | GGCTGGATTGACATGAGGAG | RT-PCR transcript detection | 59R2 |
| AtACTIN8 | ATGAAGATTAAGGTCGTGGCA | RT-PCR transcript detection | ACTIN8F1 |
| AtACTIN8 | TCCGAGTTTGAAGAGGCTAC | RT-PCR transcript detection | ACTIN8R1 |
| pBIN-pROK 2 | AGGGATTTTGCCGATTTCGGA | Genotyping | SALK-LB |
| AtCFI59 | CATAATCAGCCTCAACCACCA | Genotyping | 59F1 |
| AtCFI59 | CAGAACCACTAGCCATGTTGT | Genotyping | 59R1 |
| AtCFI59 (*atcfi59-1*, *atcfi59-2*) | AGGGATTTTGCCGATTTCGGA | Mutant site sequencing | SALK-LB |
| AtCFI68 | GCAAGCCCATTTGTTCTGGG | Genotyping | 68F1 |
| AtCFI68 | GATGGGAGAGATCAGATGGAT | Genotyping | 68F2 |
| AtCFI68 | ACCCATCACAGGTCTCATCAA | Genotyping | 68R2 |
| AtCFI68 (*atcfi68-1*, *atcfi68-2*) | ATAACGCTGCGGACATCTACA | Mutant site sequencing | GABI-LB |
| AtCFI68 (*atcfi68-3*) | AGGGATTTTGCCGATTTCGGA | Mutant site sequencing | SALK-LB |
| pAC161 | ATAACGCTGCGGACATCTACA | Genotyping | GABI-LB |
| AtCFI25a | AACCATTCCGCAGCAGCTAT | For 3′ RACE experiments | 25aF5 |

| | | | |
|---|---|---|---|
| AtCFI25a | CGTGATTTCGACTCGGTGTG | For 3′ RACE experiments | 25aF6 |
| AtCFI25b | ACTTCAAACTCCTCGCTGTTCC | For 3′ RACE experiments | 25bF4 |
| AtCFI25b | GACTTATGGGCCGATCATGTCG | For 3′ RACE experiments | 25bF5 |
| AtCFI59 | GCAGCAGAGAGACCGTGATTCT | For 3′ RACE experiments | 59F3 |
| AtCFI59 | GATCCAGGTCAAGAGATGCAGA | For 3′ RACE experiments | 59F4 |
| AtCFI68 | GATGAGGAATGGAACAGAGGCC | For 3′ RACE experiments | 68F4 |
| AtCFI68 | GATTATGGGAAAAGAAGGCGGC | For 3′ RACE experiments | 68F5 |
| AtACTIN8 | TGAAGATTAAGGTCGTGGCACC | For 3′ RACE experiments | ACTIN8F2 |
| AtACTIN8 | CAGAGTATGATGAAGCAGGTCC | For 3′ RACE experiments | ACTIN8F3 |
| general | Undisclosed. Designed by Takara to have dT region with BamHI, KpnI, and XbaI sites. | For 3′ RACE experiments | Oligo dT-3sites Adaptor Primer |
| general | CTGATCTAGAGGTACCGGATCC | For 3′ RACE experiments | 3 sites Adaptor Primer |
| AtCFI25a | AAGGAAAAAAGCGGCCGCTATGGCTATGTCTCAG GTGGTG | Cloning into pENTR vector | 25a-pENTR-F |
| AtCFI25a | TTGGCGCGCCTCGAACTAATCATGTTGAAATGGAA T | Cloning into pENTR vector | 25a-pENTR-R |
| AtCFI25a | TTGGCGCGCCTTCACGAACTAATCATGTTGAAATG | Cloning into pENTR vector | 25a-pENTR-R-w ithSTOP |
| AtCFI59 | AAGGAAAAAAGCGGCCGCTATGACTGAAGAAAAC GATTATGGAG | Cloning into pENTR vector | 59-pENTR-F |
| AtCFI59 | TTGGCGCGCCTGTCACCTCGTCTCCTCTTTCC | Cloning into pENTR vector | 59-pENTR-R |
| AtCFI59 | TTGGCGCGCCTTCAGTCACCTCGTCTCCTCTTT | Cloning into pENTR vector | 59-pENTR-R-wit hSTOP |
| AtCFI68 | AAGGAAAAAAGCGGCCGCTATGGATGAGGGAGAT GGGA | Cloning into pENTR vector | 68-pENTR-F |
| AtCFI68 | TTGGCGCGCCTTTCAGTAGTAAGCCGCCTTCTTT | Cloning into pENTR vector | 68-pENTR-R |
| AtCFI68 | TTGGCGCGCCTTCATTCAGTAGTAAGCCGCCTT | Cloning into pENTR vector | 68-pENTR-R-wit hSTOP |

| | |
|---|---|
| *AtCFI59* and *AtCFI68*, without NLS signals: Reverse primers to amplify deletion sequences of shorter versions of AtCFI59 and AtCFI68 (amino acid sequences lacking C-terminal NLS signals). The forward primers are the same set used for the amplification of the full-length nucleotide sequences of *AtCFI59* and *AtCFI68* (Cloning into pENTR vector). | |
| CFI59_ΔNLS_R_ | AAGAAAGCTGGGTCGGCGCGCCtGTGACTATCTTTCTCTTCACGGTG |
| CFI59_ΔNLS _R_STOP | AAGAAAGCTGGGTCGGCGCGCCttcaGTGACTATCTTTCTCTTCACGGTG |
| CFI68_ΔNLS _R_ | AAGAAAGCTGGGTCGGCGCGCCtGCCTCTGTTCCATTCCTCATC |
| CFI68_ΔNLS _R_STOP | AAGAAAGCTGGGTCGGCGCGCCttcaGCCTCTGTTCCATTCCTCATC |

**Supplementary Table S2.**
Primers used in this study for PAT-seq.

| | |
|---|---|
| RT_primer_universal for PAT-seq | GTTCAGAGTTCTACAGTCCGACGATCNNNNNNTTTTTTTTTTTTTTTTTTVN |
| Switch Primer for PAT-seq | CCTTGGCACCCGAGAATTCCAGGG |

**Supplementary Table S3.**
Results from the QuantifyPoly(A) tool of alternative polyadenylation dynamics across a whole gene.

The excel file contains six tables comparing *atcfi59*, *atcfi68*, *atcfi59 atcfi68*, *atcfi25a*, *atcfi25b*, *atcfi25a atcfi25b* mutants with wild-type plants. Each table contains information on the gene ID, percentage difference metric measuring the dynamics induced by changes in biological condition (pd), r indicating the lengthening (positive values) or shortening (negative values) status (r), and statistical test p value (p-value).

**Supplementary Table S4.**
Results of the co-IP experiments.

The excel file contains three tables with the results of co-IPs from GFP-AtCFI25a, GFP-AtCFI59 and AtCFI68-GFP experiments, together with the statistical analysis.

**Supplementary Table S5.**
Results of differential gene expression analysis based on negative binomial distribution (DEseq2).

The excel file contains six tables comparing *atcfi59*, *atcfi68*, *atcfi59 atcfi68*, *atcfi25a*, *atcfi25b*, *atcfi25a atcfi25b* mutants with wild-type plants (Col-0).
